# A 3D image atlas chronicling cellular and structural dynamics following lung injury identifies the aberrant expansion of endothelial cells that fail to form perfused vasculature

**DOI:** 10.64898/2026.07.22.740063

**Authors:** Jingyi Xia, Arjun Gupta, Ethan Poupard, Hayden Helms, Brendon M. Baker

**Author notes:** Corresponding Author: Brendon M. Baker, Ph.D., Assistant Professor, Department of Biomedical Engineering, University of Michigan, 2174 Lurie BME Building, 1101 Beal Avenue, Ann Arbor, MI 48109.

## Abstract

Intratracheally delivered bleomycin in mice is the most widely used in vivo model of pulmonary fibrosis, yet key aspects remain poorly defined, including sex-dependent responses and the temporal peak of injury. Standard 2D histology further overlooks regional heterogeneity and cannot resolve the 3D architecture or connectivity of endothelial cells (ECs). Here, we established a multi-scale 3D imaging pipeline integrating precision-cut lung slices from EC lineage-tracing mice, optical clearing, and AI-driven 3D segmentation to map cellular and structural dynamics from whole-lobe tile scans to single-cell resolution. We identified sex as a critical biological variable, with males exhibiting a delayed but more severe fibroproliferative response. Unsupervised K-means clustering identified three distinct tissue microenvironments: healthy parenchyma(KMC1), a myofibroblast-rich fibrotic core (KMC2), and a previously uncharacterized EC-dense perilesional region (KMC3) defined by massively expanded but non-perfused ECs that acquire a pro-inflammatory phenotype. This aberrant endothelial response precedes peak myofibroblast accumulation and persists beyond fibrotic resolution, leaving a ‘vascular scar’ that extends into the large-vessel hierarchy. Together, this 3D image atlas, made publicly available as an interactive resource [https://mosaic-lung.com/], reveals the activated endothelium as an underexplored therapeutic target in pulmonary fibrosis.

## INTRODUCTION

Fibrosis, or excessive tissue scarring, arises from an aberrant and persistent wound healing response to acute or chronic tissue injury^1,2^ and is often fatal when it occurs in vital organs like the liver, heart, or lung^3,4^. In particular, lung fibrosis is the primary determinant of fatality for numerous interstitial lung diseases (ILDs), including idiopathic pulmonary fibrosis (IPF)^1,2^, chronic hypersensitivity nonspecific interstitial pneumonia^6^, all of which involve scarring of the alveolar parenchyma resulting in impaired gas exchange and eventual respiratory failure. Viral infections have recently been identified as a significant risk factor for lung fibrosis, with studies indicating survivors of severe or prolonged COVID-19 are predisposed to lung fibrosis^7–9^. Given the vast population affected by COVID-19, lung fibrosis may emerge as a substantial global health challenge as the cohort of survivors ages.

Currently, only three FDA-approved drugs are available for treating lung fibrosis^10^. While these therapies slow or halt the fibrotic progression, they fail to reverse fibrotic changes in the lung and improve or restore lung function^11^. Successful development of more effective drugs for treating lung fibrosis has remained challenging^12^. This difficulty stems from several factors, including the complexity of pro-fibrotic signaling networks which involve other cell types beyond the (myo)fibroblast^13^, the lack of *in vivo* models that adequately capture the sequelae of human disease^12^, and likely the historical overemphasis on studying MF biology while ignoring other key stromal cell types^14^. Despite the lung’s extensive vascularization and the sheer abundance of endothelial cells (ECs), the contributions of ECs to the pathogenesis of fibrosis are not established^15^. While EC dysfunction and a loss of vascular integrity have been associated with IPF and acute respiratory distress syndrome (ARDS)^16–18^, it is unclear whether these vascular changes are merely bystander effects or if ECs actively contribute to the fibroproliferative cascade. Resolving this question is essential to building on our understanding of lung fibrosis towards identifying novel and improved therapies.

The intratracheally delivered bleomycin model remains the most widely employed *in vivo* model of lung fibrosis but our understanding of the vascular response has been historically limited by the means of analysis. Following initial EC injury and increased vascular permeability, a phase of angiogenesis is believed to occur, but the resulting neovasculature is considered aberrant and dysfunctional^19–21^. The conventional use of histological approaches to assess the vascular response has restrained progress. First, the inherently 3D architecture of vascular networks including vessel density, branching patterns, and tortuosity are all inaccessible by histology. For example, thin sections cannot distinguish whether newly emerging ECs represent true angiogenic sprouts connected to a parent vessel or isolated, disconnected ECs from an injured endothelium. Second, most previous studies have focused on hotspots of the injury response, overlooking the spatial heterogeneous injury response and vascular remodeling. Together, conventional analyses provide only structurally limited snapshots of an architecturally complex, spatially heterogeneous response. Beyond these limitations, our understanding has been further hampered by limited analysis of spatiotemporal dynamics, sex-specific responses, and vascular/EC dynamic. For instance, bleomycin-induced injury is self-resolving in young mice but partially progressive in aged mice, a nuance critical for experimental design^20,22^. Without a sex-specific spatiotemporal map defining peak response and key shifts in cellular populations, endpoint interpretations can be misguided and sex-dependent variations can reduce experimental reproducibility.

To address these gaps, here we combined EC lineage tracing, tissue clearing, and unbiased K-means clustering (KMC) of thick sections of entire lung lobes, which enabled objective classification of spatially distinct lung regions for single cell resolution high magnification imaging and AI-guided volumetric image analysis. Together, this multi-scale approach generated a spatiotemporal 3D atlas of the lung injury and repair process. Our analysis reveals an intriguing biological phenomenon which would be invisible to conventional histology: an extensive, injury-induced expansion of ECs that remain largely disconnected from systemic circulation. These findings identify the non-perfused vascular niche as a distinct feature of resolving lung injury. This raises key questions about the functional role of these disconnected ECs: do they actively promote fibrotic progression or contribute to its resolution, and could the activated endothelium serve as a therapeutic target distinct from existing myofibroblast-centric antifibrotic strategies? Our 3D imaging datasets and KMC analysis pipeline have been made publicly available as MOSAIC-Lung (https://mosaic-lung.com/), a community resource for those interested in vascular remodeling and injury-repair dynamics in the lung.

## RESULTS AND DISCUSSION

### A multi-scale 3D imaging pipeline reveals sex-dependent heterogeneity and establishes a standardized spatiotemporal atlas of lung injury

To enable lineage tracing of ECs during lung injury, we employed the Tie2-Cre/mTmG mouse model, where Tie2 promoter activity drives a permanent genetic switch from membrane-localized tdTomato to GFP expression (**Fig. 1A**). This constitutive driver is the most widely validated tool for pan-endothelial lineage tracing ^23^, broadly labeling ECs, from mature vessels through immature tip cells and nascent sprouts. This is a deliberate design choice: VE-Cadherin-based (Cdh5) drivers label only mature ECs, making them poorly suited to capture the nascent endothelial populations ^24–27^. Inducible Cdh5(PAC)-CreERT2 models further suffer from stochastic tamoxifen-dependent recombination, producing heterogeneous GFP labeling in our hands. While Tie2-Cre activity extends to hematopoietic progenitors—a limitation shared by other drivers common to endothelial-hematopoietic progenitors including the constitutive Cdh5-Cre model ^28^, we address this limitation by identifying macrophages with F4/80 immunostaining (Supplementary Fig. 4 Additionally, across our imaged FOVs, encompassing both alveolar parenchyma and airspace, macrophages were sparse (1–2 per FOV) and morphologically distinguishable from elongated ECs by their rounded shape. While macrophages were more abundant in the perivascular space around large vessels, particularly at 7 DPI (day post injury), these regions fall outside our alveolar-focused quantification framework. The Tie2-Cre/mTmG system therefore provides the most robust strategy for capturing the full endothelial compartment in the context of this study.

**Figure 1:**
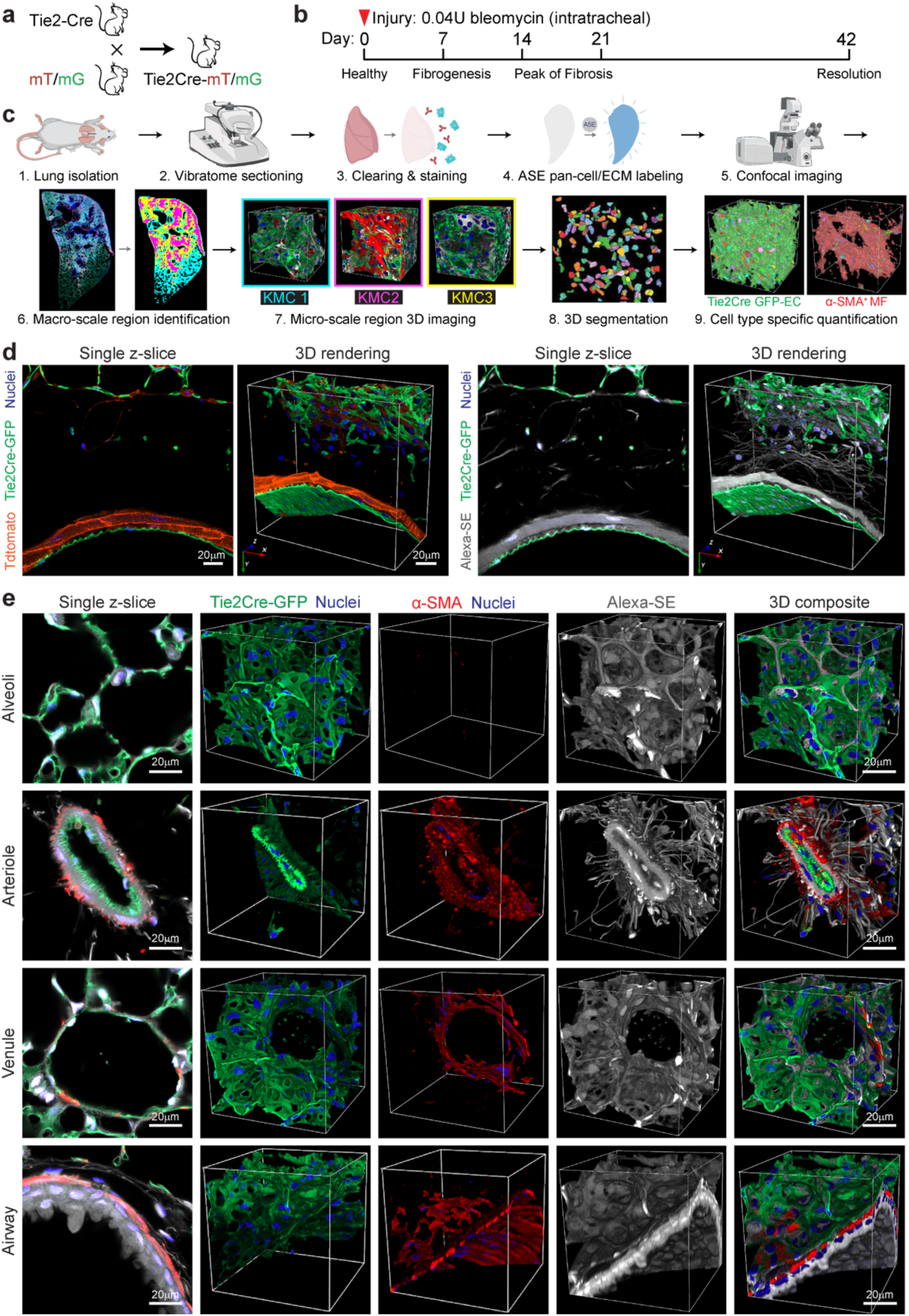
A multi-scale 3D imaging and analysis pipeline for spatiotemporal characterization of lung injury. (a) Mouse breeding strategy for EC lineage tracing. (b) Timeline for bleomycin murine lung model. (c) Pipeline workflow: Vibratome-sectioned lung slices (PCLS) undergo CUBIC clearing then pan-tissue labeling with ASE. Whole-lobe tile scans are classified via unsupervised K-means clustering (KMC) to unbiasedly identify distinct tissue microenvironments (KMC1 to 3) for guiding high-resolution confocal imaging, followed by AI-driven 3D segmentation and quantification. (d) ASE as pan tissue labeling complement Tdtomato cellular signaling: Representative images show Tie2-GFP labeling of the endothelial network against endogenous cellular tdTomato (left) versus ASE pan-tissue counterstain (right) visualize additional extracellular matrix. (e) 3D volumetric characterization of lung structures: Representative 3D renderings of alveoli, arterioles, venules, and airways in healthy control mouse lung ECs (Tie2-GFP, green), smooth muscle cells (α-SMA, red), and pan-tissue (ASE, white).

We developed a 3D visualization pipeline by integrating tissue clearing, confocal imaging, and AI-guided cell segmentation to resolve post-injury lung remodeling at single-cell resolution, with particular attention to endothelial contributions that remain poorly defined in fibrosis^15,18^. Implementing this pipeline first required standardizing the bleomycin model. We addressed previously reported sources of variability, including bleomycin dosage and time of assessment. While many studies dose based on body weight (1.0-3.0 U/kg)^30,31^, we administered a single, fixed dose (0.04 U) to all mice irrespective of weight or sex, given that lung volume is relatively constant in adult mice despite differences in sex and weight ^32^(**Fig. 2C, left and middle**). Second, we performed characterization across injury and resolution phases (7, 14, 21, and 42 days post-injury (DPI)) (**Fig. 1B),** capturing injury and fibroproliferation (days 7 and 14 DPI), established fibrosis (day 21 DPI), and spontaneous resolution (day 42 DPI)^30,33^. Supporting resolution over this range of time points, both lung collagen content and dry weight declined between 21 DPI and 42 DPI.

**Figure 2:**
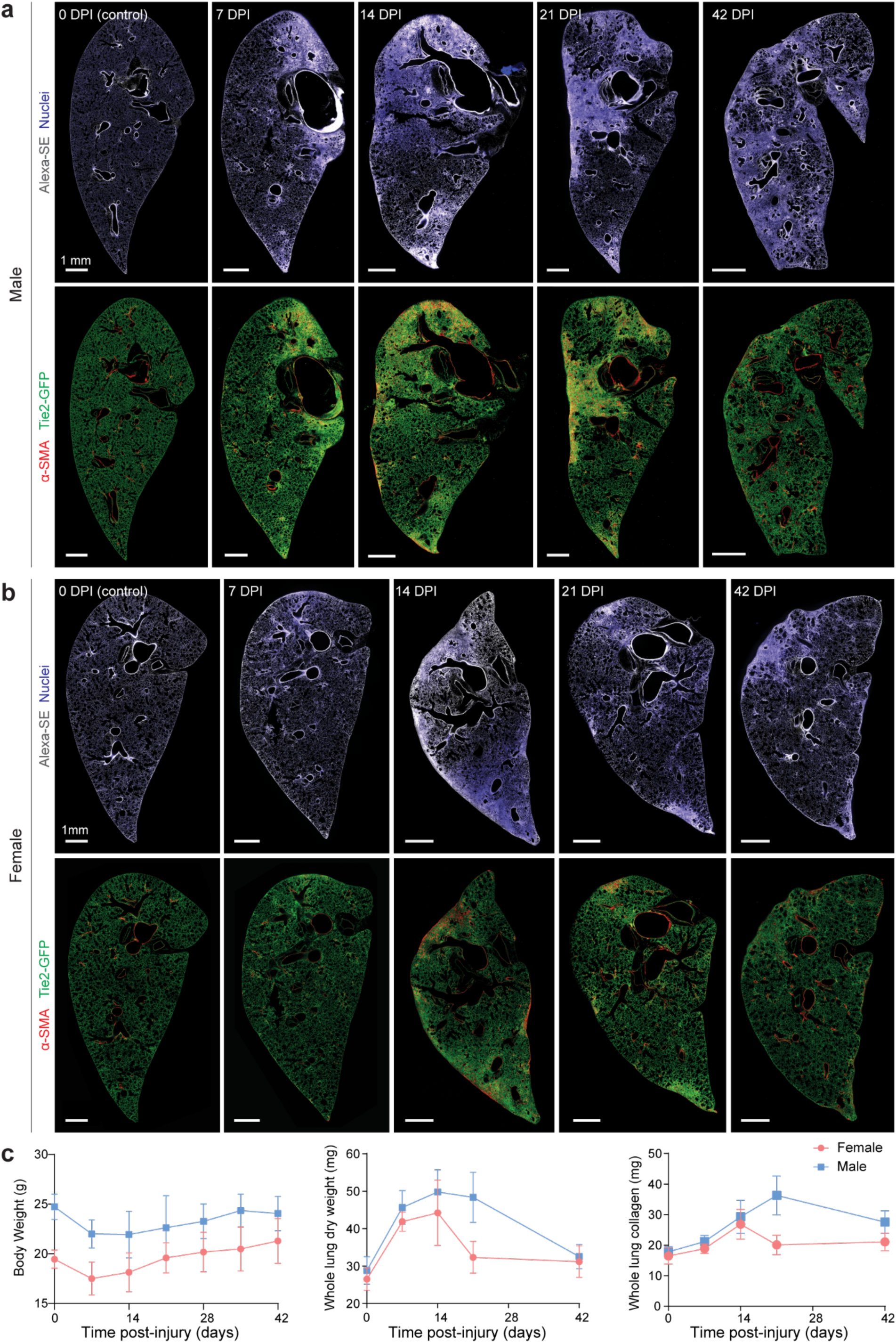
Sex-dependent spatiotemporal dynamics of bleomycin-induced lung injury. (a, b) Representative whole-lobe tile scans of male (a) and female (b) lungs across the injury time course (0 DPI–6). Top rows: tissue architecture (Alexa-SE, white) and nuclei (blue). Bottom rows: ECs (Tie2-GFP, green) and MFs (α-SMA, red). Note more severe responses in males compared to in females. (c) Bulk whole lung assessment. Body weight (left), whole lung dry weight (middle), and total lung collagen content via hydroxyproline assay (right) over study time course. Data are mean ± s.d. N >= 5 mice for both male and female.

Conventional histological assessment of lung fibrosis relies on thin (typically < 10 μm) sections, which precludes analysis of the inherently 3D architecture of the tissue and embedded vasculature. This limitation is particularly consequential for the pulmonary vasculature, whose network organization, branching patterns, and connectivity cannot be resolved from individual thin sections. To overcome this, our pipeline integrates tissue clearing of thick (200 μm) precision-cut lung slices with confocal volumetric imaging and AI-guided cell segmentation^27^. We additionally employed pan-protein labeling with Alexa Fluor succinimidyl esters (ASE) as a counterstain to visualize cell and extracellular matrix (ECM) architecture (Fig. 1D). ASE complements EC vs. non-EC lineage tracing by labeling all amine-containing proteins, including acellular ECM components (Fig. 1E).

A second historical challenge arises from the spatial heterogeneity inherent to fibrotic lung disease. A “patchy” pathology is a key feature of human IPF^2^ and the bleomycin mouse model effectively recapitulates this, presenting a mosaic of fibrotic and conserved healthy tissue. This is likely due to the heterogeneous distribution of bleomycin delivered as a liquid bolus via oropharyngeal aspiration, in contrast to intratracheal aerosolization which produces a more uniform distribution of lung injury^34^. Although whole-lobe sections can in principle be assessed by traditional histology, most studies focus on selected fibrotic “hotspots,” which introduces selection bias. To address heterogeneity in an unbiased fashion, we systematically evaluated entire lung lobe sections via tile-scanning. We then employed unsupervised k-means clustering (KMC) to identify distinct tissue regions based on Tie2Cre-driven GFP expression, α-SMA immunostaining, and nuclear density (DAPI staining). Unsupervised clustering defined a spatial map which informed the acquisition of high-magnification 3D image sets for single-cell resolution analyses (Fig. 1C).

### Sexual dimorphism in the severity and temporal kinetics of bleomycin-induced lung fibrosis

To address sex as a biological variable, we characterized the fibrotic injury response in male and female mice. Whole-lung tile scan imaging suggested a similar pathological sequence in both sexes: an initial expansion of GFP⁺ ECs at 7 DPI, followed by the appearance of α-SMA⁺ myofibroblast (MF)-rich fibrotic regions at 14 to 21 DPI, which largely resolved by 42 DPI (**Fig. 2A, B**). However, the fibrotic response was visibly less severe in females, with smaller regions of injury and overall lower α-SMA signal. This temporal characterization following lung injury informs the selection of time points for specific experimental goals. For instance, our data reveal that by 42 DPI, MF-enriched regions have largely resolved, while both deposited ECM (Fig. 2C) and an expanded EC population persist (Fig. 2A, 42 DPI). Because MFs are more transient in this model while changes in ECM and ECs are more persistent, late-endpoint readouts (e.g., 42 DPI MF counts) cannot reliably distinguish a drug-mediated decrease in MF from that arising spontaneously. This consideration supports the field’s most common focus on 21 DPI for assessing therapeutic targeting of MFs ^30,31,33^. For studies evaluating fibrosis reversal, alternative models such as aged-mouse or repetitive-dosing protocols ^35^ are therefore required, where fibrosis persists long enough to distinguish drug-induced regression from the spontaneous MF resolution our data reveal.

These qualitative observations were supported by quantitative measurements. Following lung injury, a significant increase in lung dry weight occurred at 7 DPI, preceding a rise in total lung collagen content. This suggests an initial phase of cellular infiltration — consistent with the early inflammatory response and the EC expansion noted above — prior to overt fibrotic ECM deposition. Subsequently, total collagen content, measured by hydroxyproline (OHP) assay, increased markedly. The fibrotic response was attenuated in females and peaked earlier (14 DPI) compared to males (21 DPI), with total lung collagen beginning to decline by 21 DPI (**Fig. 2C, middle and right; online atlas**). By 42 DPI, lung dry weight and OHP levels declined toward baseline in both sexes, consistent with resolution of the fibrotic injury response. Notably, this attenuated and temporally shifted response in females occurred despite females receiving a higher effective dose relative to body weight, underscoring sex-specific differences in the injury response. These findings align with the sexual dimorphism reported in human fibrotic diseases, where males exhibit a higher incidence and greater severity of lung fibrosis^24–26^. Together, our data establish sex as a critical biological variable in the bleomycin model. To preserve resolution of these sex-dependent temporal kinetics, we recommend that future studies analyze male and female cohorts separately rather than pooling the data.

To probe systemic drivers of the EC expansion (Fig. 2A, B), we profiled circulating angiogenic and inflammatory factors in serum across the injury time course. Due to a limited number of samples and given the cost of analysis, we assessed male mice only; whether female mice exhibit the same systemic kinetics remains to be determined. A robust pro-angiogenic program was established as early as 7 DPI and persisted through 21 DPI (Supplementary Fig. 2a), characterized by sustained upregulation of endoglin — a TGFβ co-receptor that signals through ALK1 (ACVRL1) to activate Smad1/5/8 angiogenic sprouting — and PDGF-AA (Supplementary Fig. 2b). Acute inflammatory markers (IL-1α, Coagulation Factor III) showed a transient spike at 7 DPI before declining, while the mitogenic and matrix-preserving drivers TIMP-1 and PDGF-AA remained elevated, consistent with the sustained EC expansion observed by imaging.

### Unsupervised spatial clustering identifies tissue regions composed of proliferative and inflammatory endothelial cells

To objectively classify tissue microenvironments throughout the fibroproliferative and resolution phases of lung injury, we applied unsupervised K-means clustering (KMC) to whole lobe tile-scan images of DAPI-stained nuclei, Tie2Cre-GFP, and immunostained αSMA. We focused this analysis on males given the more severe fibrotic responses noted in the studies above (**Fig. 2**). The elbow method was used to determine the optimal cluster number (K). The resulting plots indicated a clear inflection point between K=2 and K=3 (**Supplementary Fig. 1a, b**). K=2 broadly segregated the lung into ‘healthy’ and ‘diseased’ regions. However, inspection of the feature space revealed significant biological heterogeneity remained within the single ‘diseased’ cluster **(Supplementary Fig. 1c)**. Specifically, this cluster was composed of sub-populations dominated by either MFs (α-SMA^+^) or ECs (Tie2-GFP^+^). Consequently, we selected K=3 for all bleomycin-treated groups to capture this level of granularity. For uninjured control lungs (0 DPI), two clusters were sufficient to distinguish healthy parenchyma from large structural features such as large smooth muscle covering vasculatures and airways.

The K=3 model partitioned the tissue into three biologically distinct microenvironments (**Fig. 3a**). KMC1 corresponded to healthy parenchyma, conserved over time and characterized by low nuclear density and minimal α-SMA or GFP signal. In contrast, KMC2 represented the classic fibrotic lesion arising from lung injury, defined by heightened α-SMA intensity and nuclear density reflecting the accumulation of myofibroblasts. Notably, KMC3 corresponded to a previously uncharacterized perilesional microenvironment with high Tie2Cre-GFP signal and lower but present α-SMA expression. To visualize the temporal dynamics of each region, we projected cluster centroid trajectories onto the Tie2Cre-GFP vs. α-SMA intensity axes (**Fig. 3b, c**). The clear separation between centroids confirmed that these clusters represent distinct biological entities.

**Figure 3:**
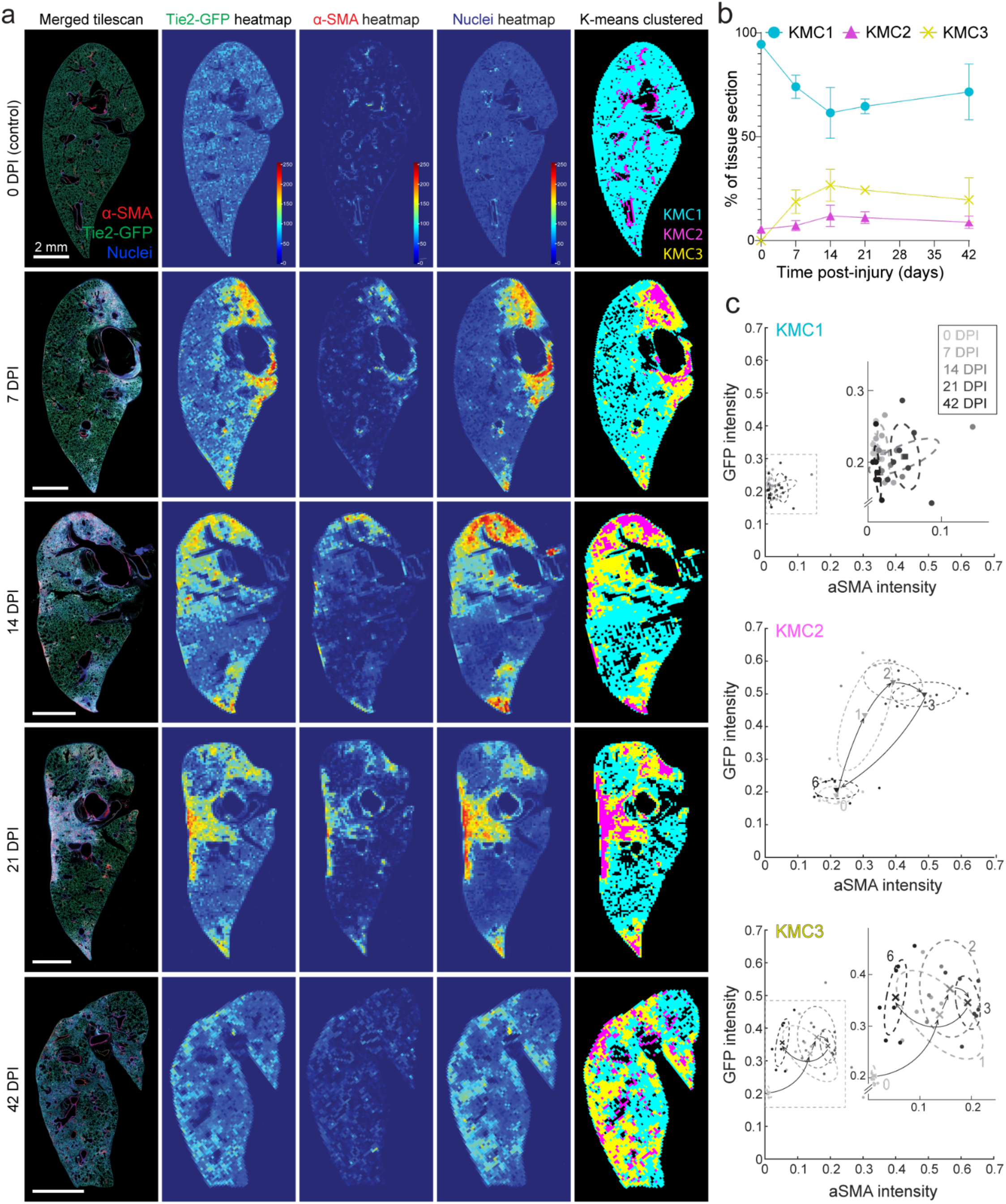
Unsupervised K-means clustering identifies distinct spatial microenvironments conserved across different phase of lung injury. (a) Representative time-course whole-lobe multichannel tile scans (left) with corresponding single-channel intensity heatmaps (Tie2-GFP, α-SMA, Nuclei) (middle), and K-means clustered maps (right). Clustering was performed on binned Tie2-GFP, α-SMA, Nuclei intensity spatial data to segment tissue into three distinct phenotypes: KMC1 (cyan, healthy), KMC2 (magenta, MF-dense core), and KMC3 (yellow, EC-dense perilesional region). (b) Quantification of the tissue area fraction occupied by each cluster. Data are mean ± s.d. N>=3 mice per time point and n>2 tile scans per mouse. **(c)** Temporal trajectories of cluster centroids. Centroids were calculated in 3D space (GFP, α-SMA, DAPI) and projected onto the Tie2-GFP vs α-SMA feature space to visualize the divergence between endothelial and myofibroblast phenotypes. Note the distinct locations and trajectory of the KMC3 regions (high GFP, low α-SMA) compared to the classic fibrotic KMC2 regions (moderate GFP, maximal α-SMA).

KMC1 remained stable over time, while KMC2and KMC3 displayed distinct trajectories. At 7 DPI, Tie2Cre-GFP intensity rose sharply in both KMC2 and KMC3, while α-SMA in KMC2 was just beginning to increase. From 14-21 DPI, KMC2 retained elevated GFP intensity and increasing α-SMA as the fibrotic lesion matured; KMC3 retained its high-GFP, low α-SMA signature at these time points. By 42 DPI, KMC2 returned to baseline (suggesting resolution of fibrosis), but KMC3’s GFP intensity remained elevated, indicative of a persistent increased in ECs after the fibrotic core resolved.

We next performed high-magnification imaging to resolve the cellular composition and architecture of each KMC region at single-cell resolution and to visualize and quantify the cellular dynamics underlying centroid trajectories (Fig. 4a). KMC1 regions retained their normal alveolar architecture and low cell density across the entire study period, confirming the spatial heterogeneity of the injury response. In KMC2, we observed a marked loss of airspace, a 4- to 5-fold expansion of Tie2Cre-GFP^+^ ECs starting at 7 DPI, and an accumulation of α-SMA^+^ MFs that peaked at 21 DPI (Fig. 4c). The perilesional KMC3 microenvironment exhibited a distinct 3D phenotype: dense, complex tubular networks of Tie2Cre-GFP^+^ ECs encased the fibrotic core, while the rare MFs present formed loose, mesh-like α-SMA arrays rather than the thick stress fibers seen in KMC2 (Fig. 4a). Quantitatively, this produced markedly different EC-to-MF ratios at peak fibrosis (21 DPI) — 2:1 in KMC2 versus 7:1 in KMC3 — and EC counts in both regions remained elevated through 42 DPI even as MFs resolved in KMC2 and 3 (Fig. 4c).

**Figure 4:**
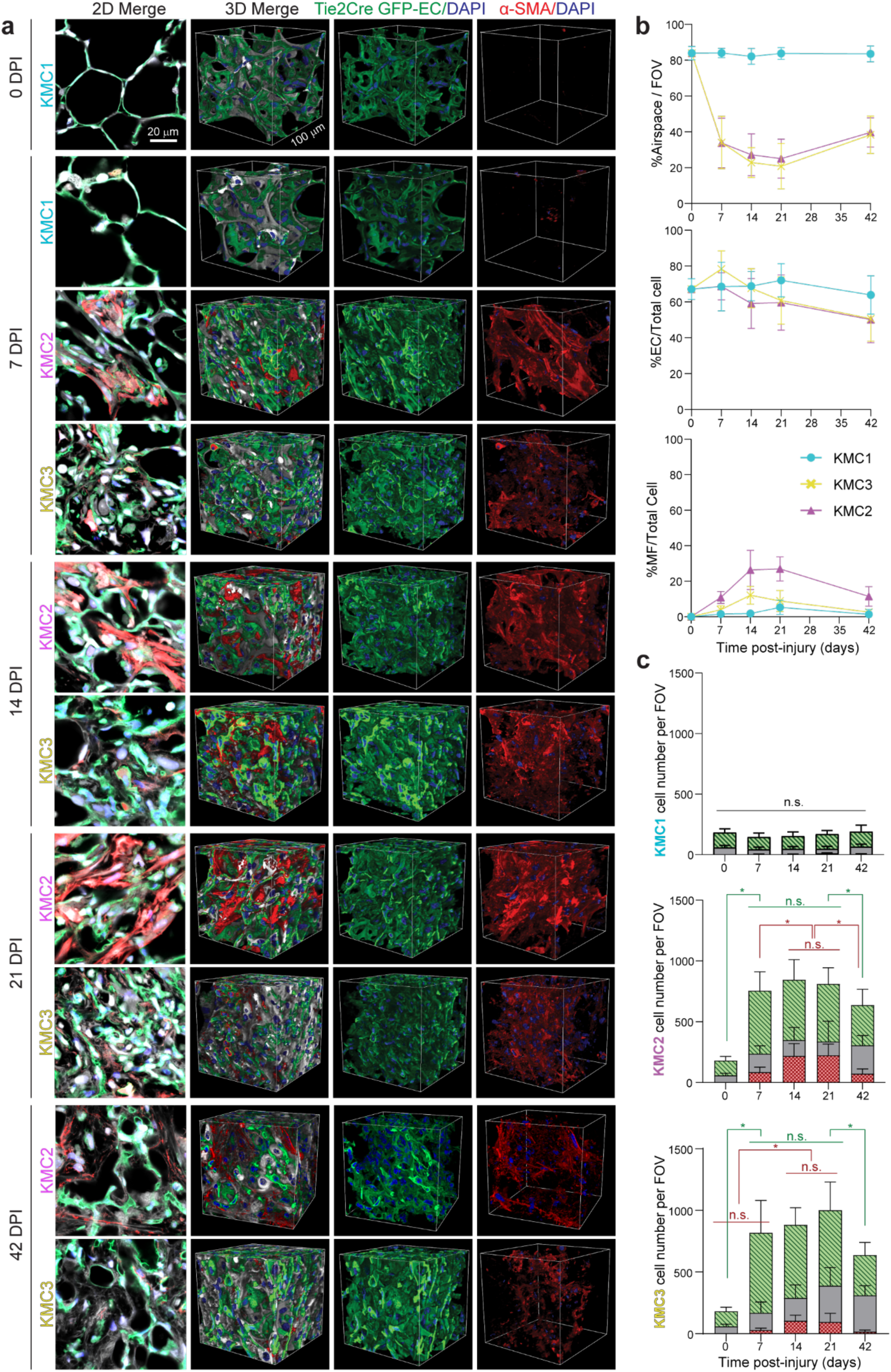
High-resolution 3D characterization reveals a distinct endothelial-dense, myofibroblast-sparse microenvironment (KMC3). **(a)** Representative high-magnification 2D slices and 3D volumetric renderings of KMC-classified regions (KMC1–3) across lung injury timeline. KMC1 retains healthy alveolar architecture. KMC2 fibrotic core exhibits dense accumulation of both MFs (α-SMA, red) and ECs (Tie2-GFP, green). In contrast, the perilesional KMC3 region exhibits pronounced EC densification with minimal MFs. **(b)** Quantification of airspace preservation and cell-type proportions to total cell count. **(c)** Absolute cell counts per field of view (FOV) by cell type. At peak fibrosis (21 DPI), KMC3 is defined by a marked ratio of ECs to MFs, distinct from the mixed cellularity of the KMC2 fibrotic core. Data are mean ± s.d. N >= 3 mice per time point and n >= 5 high mag 3D image per KMC regions per mouse. 3D image FOV volume is ∼ 10^6^ μm^3^.

Sustained endothelial expansion in KMC3 could in principle arise from local EC proliferation or recruitment of endothelial progenitor cells (EPCs) from circulation. While EPC recruitment has been historically proposed, recent lineage-tracing studies in lung and other organs have found limited evidence for bone marrow-derived contribution to adult endothelial repair, instead implicating local EC proliferation as the dominant mechanism^36–38^. To test whether local proliferation underlies KMC3 expansion, we examined Ki67 immunostaining across the injury time course. At baseline (week 0), the lung vasculature was quiescent, with virtually no Ki67^+^ GFP^+^ nuclei detected (**Supplementary Fig. 3a (left), 3b**). However, 7 DPI marked a peak in EC proliferation, with Ki67^+^ GFP^+^ nuclei enriched in emerging KMC3 regions, some of which were dual-positive (Tie2-GFP^+^/α-SMA^+^) cells. Morphologically, these proliferating GFP^+^ cells appeared elongated and aligned, distinguishing them from the rounded appearance of resident or infiltrating leukocytes (**Supplementary Fig. 3c**), although definitive lineage tracing would be required to rule out all immune subsets. These elongated cells were arranged in fragmented linear tracks suggestive of disconnected ECs attempting vessel formation, rather than forming the continuous networks of healthy vasculature. This localized proliferation appears to drive the formation of the dense, disorganized KMC3 microenvironment—representing an abnormal vascular expansion. By week 2, the proliferative landscape shifted. While EC proliferation persisted, the overall frequency of Ki67^+^ ECs decreased compared to 7 DPI (**Supplementary Fig. 3d**). Concurrently, we observed Ki67^+^/αSMA^+^ proliferating myofibroblasts within the fibrotic core (KMC2), consistent with recent reports that αSMA^+^ myofibroblasts retain proliferative capacity during tissue repair^39,40^. This in situ expansion, potentially driven by elevated PDGF-AA, contributes to densification of the fibrotic lesion.

To probe the inflammatory state of this expanded endothelial population, we next assessed ICAM1 and VCAM1, canonical NF-κB-driven adhesion molecules that mark endothelial inflammation in response to disturbed flow or disrupted cell-cell junctions^41–43^, and observed kinetic profiles distinct in both timing and spatial distribution. ^4445^ VCAM1 expression localized to discrete tissue compartments that changed between regions across the injury time course. Virtually absent at baseline (0 DPI), VCAM1 intensity increased markedly by 7 DPI, but was confined to the endothelium of large vessels, with minimal expression in the alveolar capillaries (**Supplementary Fig. 5a**). High-magnification imaging (**Supplementary Fig. 5b**) revealed a marked morphological shift: VCAM1^+^ ECs lining the inner arteriole wall lost the smooth, flat, spread morphology of quiescent endothelium, instead appearing hypertrophic with irregular, lobulated cell bodies bulging into the vascular lumen. In addition, small, round Tie2-GFP^+^ cells were observed in the perivascular space surrounding these VCAM-1-high vessels, separated from the endothelium by a layer of smooth muscle cells. Their round morphology and the known Tie2-Cre labeling of hematopoietic progenitors suggest these are perivascular immune cells, although definitive identification would require immune-specific markers. Notably, a subset of these luminal VCAM1^+^ cells co-expressed α-SMA (dual-positive). The localization of these cells to the inner vessel wall suggests they are transitioning ECs acquiring mesenchymal traits. By 21 DPI, the distinct vascular signal dissipated into a diffuse parenchymal signal (**Supplementary Fig. 5c**), likely reflecting VCAM1 upregulation on the expanding myofibroblast population (**Supplementary Fig. 5e**), before largely resolving by 42 DPI (**Supplementary Fig. 5d**).

In contrast, ICAM1 displayed a broader, pan-pulmonary response. From a dim basal signal at 0 DPI, ICAM1 expression increased globally at 7 DPI **(Supplementary Fig. 6a)**, appearing in even KMC1 (healthy) regions **(Supplementary Fig. 6b)**. The signal continued to rise, peaking at 21 DPI within KMC2 and KMC3. Although this 21 DPI peak coincides with peak EC counts, the diffuse staining pattern and disproportionately high KMC2 signal **(Supplementary Fig. 6c, e)**, corroborated by 3D high-magnification imaging, point to substantial epithelial contribution. Although the intensity subsided by 42 DPI **(Supplementary Fig. 6d)**, it remained elevated above baseline, indicating residual endothelial activation that persists into the resolution phase. Collectively, these findings establish KMC3 as a previously uncharacterized, proliferative, and activated endothelial microenvironment that emerges early after injury and persists beyond myofibroblast resolution. Together, the disorganized, non-capillary architecture and sustained ICAM1 expression in KMC3 point to loss of EC-EC contacts as a driver of its persistent inflammatory state. Because VCAM1 can also be expressed by activated mesenchymal cells^44^ and ICAM1 by injured epithelium^45^, we restricted EC-specific interpretation to vessel-localized signals; the diffuse parenchymal signals observed at later time points (e.g., 21 DPI ICAM1 in KMC2 core) were attributed to non-EC sources.

### Injury-induced endothelial expansion drives aberrant, non-perfused vascular networks and persistent remodeling of the perivascular niche

Having shown that KMC3 endothelium is structurally disorganized and inflammatory, we next asked whether it forms functional, perfused vasculature. We labeled perfused vessels by systemic DyLight 649-Tomato Lectin (TL) injection and compared them to the full Tie2-Cre-mTmG-labeled endothelial pool (**Fig. 5A**). In healthy KMC1 regions, the vascular network displayed near-total overlap between lineage-traced and perfused vessels. Injured regions, however, exhibited a marked functional deficit: although the number of perfused ECs roughly doubled, the total Tie2-GFP^+^ population expanded far more, dropping the perfused fraction to ∼40–50% at 7 DPI and keeping it suppressed through 42 DPI (**Fig. 5B**). This perfusion deficit was evident across the entire injured lobe (Supplementary Fig. 9).

**Figure 5:**
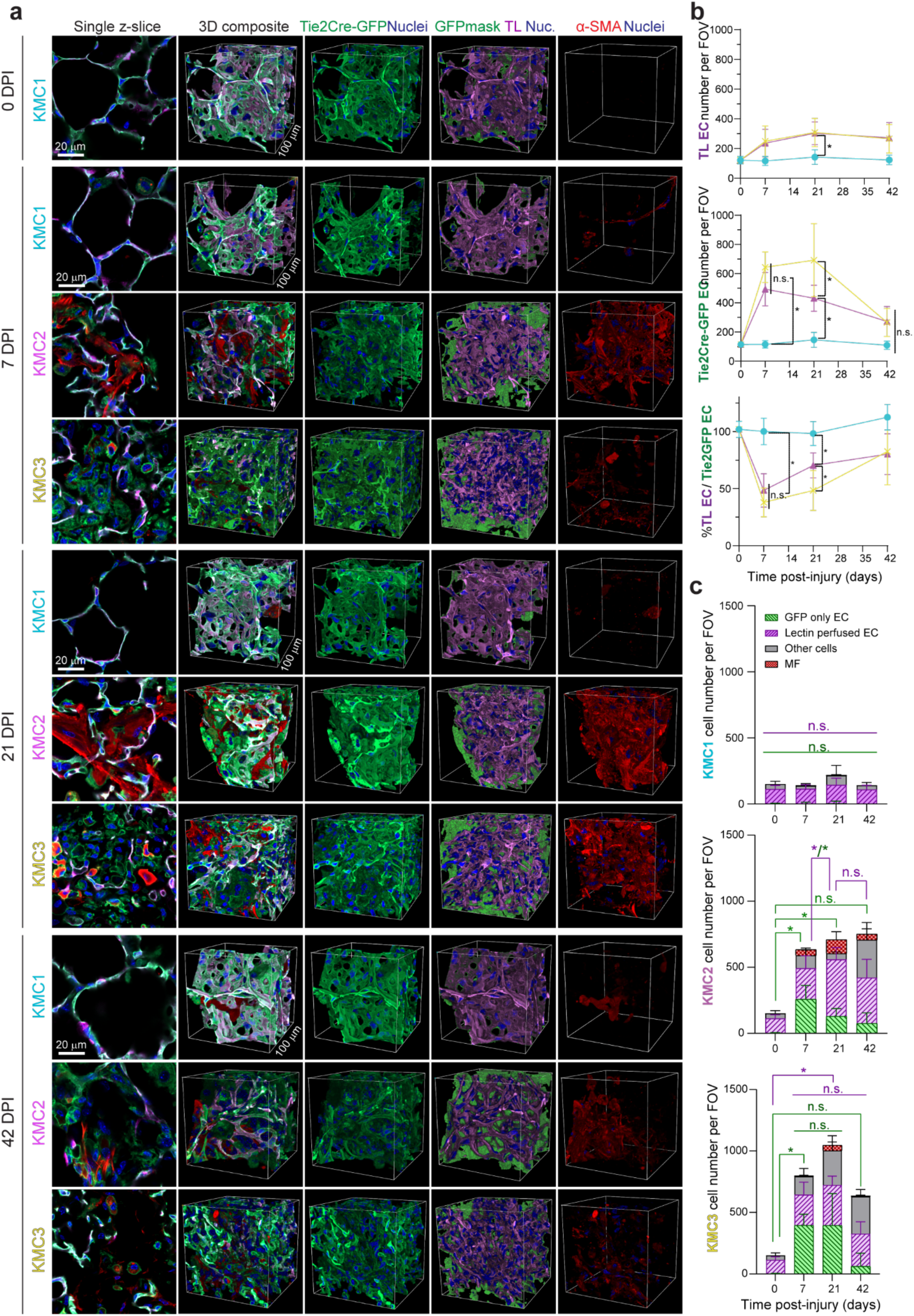
Functional assessment reveals a disconnect between endothelial expansion and vascular perfusion. (a) Representative 2D single slices and 3D volumetric renderings of KMC regions following systemic DyLight-649 Tomato Lectin (TL) injection. The Tie2-Cre-mTmG lineage trace (green) visualizes all ECs, while TL (purple) identifies mature, perfused vessels. MFs are labeled with α-SMA (red). In KMC1 regions, GFP and TL signals completely overlap, in contrast to the extensive non-perfused (GFP^+^/TL-) endothelial networks in injured KMC2 and KMC3 regions. b) Quantification of vascular dynamics per FOV. Plots display number of perfused TL^+^ ECs (top), total number of Tie2-GFP^+^ ECs (middle), and perfusion efficiency calculated as the ratio of TL^+^ to Tie2-GFP^+^ cells (bottom). (c) Stacked bar graphs detailing the absolute cellular composition of each KMC microenvironment over time. Cell population is segmented into perfused ECs (purple), non-perfused “GFP-only” ECs (green hatched), myofibroblasts (red), and other cells (grey). KMC3 is uniquely characterized by a persistent, dominant population of non-perfused ECs starting in the early phase and lasting through the peak of lung injury. Data are mean ± s.d. Significance (P < 0.05) indicates differences between KMC regions at specific time points determined by two-way ANOVA with Tukey’s multiple comparisons test. 3D image FOV volume is ∼ 10^6^ μm^3^.

Analysis of specific subpopulations revealed distinct maturation kinetics between the fibrotic core and the perilesional KMC3 microenvironment (**Fig. 5C**). In the KMC2 core, the non-perfused (’GFP-only’) population peaked at 7 DPI but declined by nearly 50% by 21 DPI. Conversely, the perfused (’Lectin^+^’) population in KMC2 doubled over the same period and remained stable through 42 DPI. This inverse relationship — non-perfused cells declining as perfused vessels increased — suggests maturation of immature sprouts into functional microvasculature, even as overall EC density remained constant. In contrast, endothelial dynamics in the perilesional KMC3 microenvironment indicated stalled angiogenesis. Here, the ‘GFP-only’ population remained dominant through 21 DPI. By 42 DPI, this non-perfused population regressed to baseline, while the fraction of perfused vessels remained elevated. Together, KMC2 sprouts appear to mature into functional microvasculature, while the majority of KMC3 sprouts remain transient and non-functional before being pruned.

These divergent vascular trajectories offer a potential explanation for the distinct pathological outcomes of these regions. The establishment of perfusion in KMC2 coincides with peak myofibroblast accumulation, suggesting that vascular maturation may be a prerequisite to support the metabolic demands of the dense fibrotic core. By contrast, the failure to establish functional perfusion in the perilesional KMC3 microenvironment coincides with the absence of a stable myofibroblast population, consistent with vascular support being a prerequisite for sustained MF activation.

The accumulation of non-perfused GFP-positive cells raises the question of cellular identity: does this population represent inflammatory infiltration (e.g., Tie2-expressing immune cells) or aberrant angiogenesis? Visual inspection supports an endothelial identity: rather than appearing as discrete, individual cells typical of leukocyte infiltration^46^, the non-perfused GFP^+^ cells appeared as dense, disorganized aggregates maintaining physical continuity with the vasculature^47^. While these cells formed interconnected clusters, they frequently lacked patent lumens or exhibited collapsed morphology, explaining the lack of lectin perfusion. This appearance is consistent with nascent tip cells and immature angiogenic sprouts that have not yet undergone lumenization.

To verify this identity, we performed targeted immunophenotyping to rule out macrophages (F4/80) and confirm a blood-endothelial origin (von Willebrand Factor, vWF) (**Supplementary Fig. 4**). F4/80 staining confirmed that macrophages were predominantly located as single cells in the adventitia, distinct from the dense parenchymal clusters we quantified. To positively confirm a blood endothelial identity, we assessed von Willebrand Factor (vWF). At baseline (0 DPI), vWF expression was low and strictly confined to the endothelium of large vessels, with virtually no signal in the alveolar capillaries (**Supplementary Fig. 7a, b**). However, paralleling the expansion of the KMC3 region, vWF signal intensified at 7 DPI and, by 21 DPI, extended aberrantly into the alveolar capillaries (**Supplementary Fig. 7c**). This abnormal presence of vWF in the distal capillary network confirms that the dense Tie2-GFP regions are ECs, albeit in a pathologically activated state.

Finally, we examined the ‘transitioning’ dual-positive cells (Tie2-GFP^+^/α-SMA+), which we hypothesized might represent an EndoMT-like phenotype. We probed these cells for E-selectin, an adhesion molecule largely restricted to activated endothelium. Unlike the broad expression of ICAM1, E-selectin signal was sparse at the whole-lobe level (**Supplementary Fig. 8a**). High-magnification imaging revealed that while baseline expression was negligible, during the acute activation phase at 7 DPI, vessels displayed intense luminal E-selectin staining, accompanied by small, round Tie2-GFP^+^ cells in the surrounding perivascular space (**Supplementary Fig. 8c**), a pattern similar to that observed for VCAM1. At the peak of fibrosis (21 DPI), the strongest E-selectin signal was mapped specifically to the dual-positive (Tie2-GFP^+^/α-SMA+) cells. High-resolution imaging revealed prominent intracellular accumulation of E-selectin in addition to surface localization. This extensive intracellular pool indicates a high biosynthetic rate and rapid membrane turnover, consistent with the sustained, hyper-activated state of ECs undergoing phenotypic transition in the inflammatory milieu. By 42 DPI, in the resolution phase, the coherent vascular signal dissipated into fragmented, punctate debris (**Supplementary Fig. 8e**). This morphological pattern is characteristic of vascular regression^27^, a resolution-phase process involving endothelial apoptosis and the proteolytic shedding of surface adhesion molecules like E-selectin^48,49^ as the excess vasculature is pruned. Together, these data establish endothelial origin for the non-perfused population in the alveolar parenchyma. We next extended our analysis to the perivascular compartment surrounding large vessels.

While our quantitative pipeline focused on the alveolar parenchyma, previous studies suggest that the adventitial and perivascular compartments serve as critical niches for fibrotic progenitors^49,50^—regions often overlooked in standard histological analyses. Leveraging our whole-lobe tile scans, we extended our qualitative analysis to these macro-structural features (**Fig. 6**). In uninjured lungs, large vessels exhibited a quiescent smooth muscle layer and a sparse perivascular space. Specifically, healthy arterioles were surrounded by a clear, largely acellular perivascular niche, while healthy venules were embedded directly within open alveoli.

**Figure 6:**
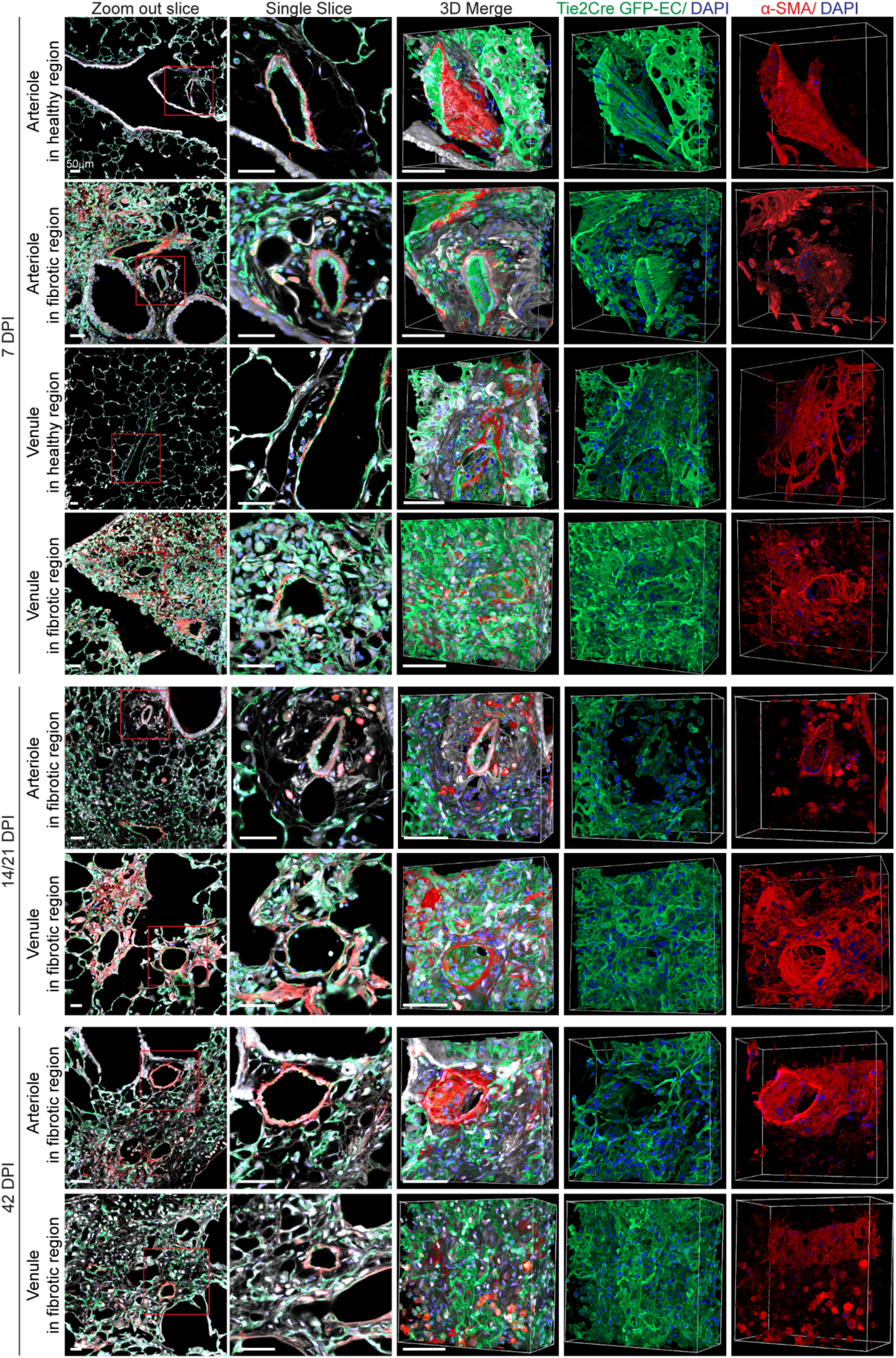
Distinct perivascular remodeling of arterioles and venules following lung injury. Representative 3D volumetric renderings and cross-sections of arterioles and venules in healthy versus fibrotic regions. In uninjured tissue (0 DPI), both vessel types maintain a quiescent phenotype with sparse perivascular cells. Following injury, the perivascular microenvironment undergoes significant remodeling. Arterioles exhibit adventitial expansion characterized by fibrillar densification and the appearance of disconnected ECs and dual-positive (Tie2-GFP^+^/α-SMA+) cells. Similarly, venules in diseased regions show a loss of the surrounding alveolar architecture, which is replaced by a dense cellular cuff containing myofibroblasts, expanded endothelial networks, and dual-positive cells. This remodeling carries into the resolution phase (42 DPI), indicating a lasting structural alteration of large vessels.

Following bleomycin injury, these regions underwent distinct remodeling trajectories. In the arteriolar niche, the previously distinct perivascular space became progressively occupied by densely packed collagen fibers. At 7 DPI, this infiltrate was characterized by the appearance of single, disconnected Tie2-GFP^+^ ECs. By 21 DPI, cellularity peaked with a dense accumulation of α-SMA+ cells and a subpopulation of dual-positive (Tie2-GFP^+^/α-SMA^+^) cells. This accumulation aligns with mechanical models where increased fiber density destabilizes endothelial VE-cadherin junctions, potentially promoting the tip-cell formation and sprouting phenotype we observed (Xia et al. 2026).

Venules exhibited a delayed but robust transformation. At 7 DPI, we observed only a few α-SMA^+^ cells surrounding the vessel. However, by 21 DPI, this evolved into a dense cellular plaque containing mature myofibroblasts and expanded endothelial networks, effectively deforming surrounding alveoli. By 42 DPI, while parenchymal fibrosis had largely resolved, the large vessel microenvironment retained a complex, expanded endothelial population. Notably, the composition of the perivenular space shifted: the mature myofibroblast population observed at 21 DPI largely disappeared, leaving behind a predominant population of dual GFP^+^ α-SMA^+^ cells. Given that these cells uniquely express E-selectin (**Supplementary Fig. 8**), their persistence during the resolution phase suggests a potential role in vascular repair or immune regulation rather than active fibrosis^51,52^. However, due to intrinsic heterogeneity in vessel diameter, branching angles, and vessel orientation throughout the lung, standardized volumetric quantification of these perivascular changes was not feasible. Nonetheless, these qualitative observations suggest that injury results in a lasting structural change to vascular hierarchy, extending beyond more transient changes in the parenchyme.

## CONCLUSION

Here, we generated a multi-scale 3D image atlas of the murine bleomycin lung injury model that captures architectural and cellular dynamics that are difficult to resolve by conventional 2D histology. This approach revealed important spatiotemporal features of the injury response, including sex-dependent differences in fibrotic kinetics, regionally distinct tissue microenvironments, and a prominent vascular remodeling program that persists beyond the resolution of canonical myofibroblast-rich fibrosis. Notably, males exhibited a delayed but more severe fibroproliferative response than females, highlighting sex as a critical biological variable in this model. These findings support sex-segregated analysis in future bleomycin studies, as pooling male and female cohorts may obscure temporally distinct injury and repair trajectories.

Through unbiased spatial clustering of whole-lobe tile-scan images, we identified an EC-dense perilesional microenvironment, KMC3, that is spatially and biologically distinct from the classic myofibroblast-rich fibrotic core, KMC2. Whereas KMC2 reflected the expected accumulation of MFs and ECM densification associated with fibrotic injury, KMC3 was defined by a pronounced expansion of Tie2-GFP+ endothelial cells with comparatively sparse MF content. High-resolution volumetric imaging and lectin perfusion further revealed that this EC expansion does not correspond to restoration of a functional vascular bed. Instead, injured regions contained dense populations of non-perfused endothelial cells arranged in fragmented and disorganized networks, consistent with immature or stalled angiogenic structures that fail to establish effective perfusion.

Temporally, this aberrant endothelial response emerged early after injury, preceding peak myofibroblast accumulation, and persisted into the resolution phase after α-SMA^+^ myofibroblast-rich regions had largely regressed. The expanded endothelial compartment was also associated with proliferative and inflammatory features, including endothelial proliferation within KMC3 and sustained expression of vascular inflammatory markers. Together, these findings suggest that the vascular response to bleomycin injury is not simply a reparative angiogenic program that restores tissue homeostasis, but instead includes a maladaptive endothelial state marked by poor perfusion, inflammatory activation, and incomplete remodeling.

The persistence of this altered endothelial and perivascular architecture reframes how resolution is interpreted in the bleomycin model. Although bulk collagen content and myofibroblast abundance decline by later time points, the injured lung does not fully return to its baseline vascular organization. Rather, the tissue retains a residual “vascular scar” characterized by persistent endothelial expansion, incomplete vascular normalization, and remodeling of the large-vessel and perivascular niches. This observation has important implications for therapeutic assessment: endpoints focused only on collagen deposition, tissue cellularity, or MF abundance may overlook persistent vascular pathology or misclassify partial repair as complete resolution.

More broadly, our findings position the activated, non-perfused endothelium as an underexplored component of lung injury and repair, and as a potential therapeutic target distinct from canonical antifibrotic strategies centered on MFs. Future studies should define the mechanisms that determine whether injury-induced EC expansion resolves into functional vasculature, regresses, or persists as a chronically activated vascular niche. In particular, cell-type-specific functional readouts, longitudinal analysis of vascular perfusion, and perturbation studies targeting EC activation, junctional stability, and angiogenic maturation will be essential to determine whether this vascular state actively contributes to fibrotic progression, impairs repair, or promotes susceptibility to recurrent injury.

*To facilitate* future *discovery, we established MOSAIC-Lung, a publicly* accessible database and image-viewing *platform containing the complete image library from this study. By enabling interactive exploration of whole-lobe architecture, regional tissue microenvironments, and high-resolution 3D vascular phenotypes, this resource provides both a foundation for investigating vascular remodeling in lung fibrosis and a framework for identifying therapeutic strategies* that promote not only MF resolution, but **true restoration of vascular and tissue architecture**.”

## ACKNOWLEDGEMENTS

JX acknowledges financial support from Rackham Predoctoral Fellowship. BMB acknowledges support from a Boehringer Ingelheim International GmbH Discovery Award, the National Scleroderma Foundation Debra Lurvey Memorial Research Grant, and a Boehringer Ingelheim International GmbH Collaborative Agreement.

## MATERIALS AND METHODS

### Murine bleomycin-induced lung injury model

B6.Cg-Tg(Tek-cre)1Ywa/J transgenic mice were crossed with Gt(ROSA)26Sortm4(ACTB-tdTomato,-EGFP)Luo reporter mice to generate progeny where Tie2-driven Cre activity resulted in permanent membrane-localized GFP expression, while non-Tie2 expressing cells retained membrane-localized tdTomato expression. Male and female mice (8-12 weeks old) were administered a single dose of bleomycin (0.04 U in saline) via oropharyngeal aspiration. Lungs and blood from injured mice were harvested at 7, 14, 21, or 42 days post-injury; uninjured mice received saline via the same route of administration and were harvested at 21 DPI as controls. For tissue collection, blood (∼0.8 mL) was collected via cardiac puncture, followed by perfusion with warm PBS until the liver blanched. Lungs were then inflated with 1.5 mL of 2% (w/v) low-melting-point agarose in 4% (w/v) paraformaldehyde (PFA). Whole lungs were extracted, fixed overnight in 4% PFA at 4°C, washed 3x with PBS, and stored in PBS with 0.01% w/v sodium azide at 4°C. Lungs designated for hydroxyproline analysis were not agarose inflated but instead flash-frozen and stored at −80°C prior to processing.

### Orthohydroxyproline collagen assay

Harvested lung tissue was lyophilized and dry weights were recorded. Tissues were papain-digested, and collagen content was determined by the hydroxyproline assay, as previously described^53^. Briefly, digests were hydrolyzed in 6N HCl at 110°C for 16 hours. Dried hydrolyzed tissue were resuspended in an acetate-citrate buffer. Samples and standards (150 µL) were added to a 96-well plate, followed by 75 µL of chloramine T solution for 20 minutes at room temperature. After adding 75 µL of p-dimethylaminobenzaldehyde solution, the plate was incubated at 70°C for 20 minutes. Absorbance was read at 540 nm after cooling. Hydroxyproline content was converted to collagen using a factor of 7.14.

### Lung tissue sectioning and clearing

Mouse whole lungs were first separated into right versus left lobes, mounted to maintain anatomical orientation, and embedded in 2% agarose for structural support during vibratome sectioning. Tissues were sectioned at 200 µm thickness and sections were stored in PBS with 0.01% w/v sodium azide. Sections were subsequently cleared using the advanced CUBIC method^54^. Briefly, sections were incubated for one day in 50% CUBIC-L, followed by one day in 100% CUBIC-L. Tissues were then blocked in PBS solution with 6 w/v% goat serum prior to immunostaining.

### Lectin perfusion and immunostaining

To label perfusable vasculature, mice were injected with 100 µL of DyLight 649-labeled Tomato Lectin (Vector Laboratories) via the right ventricle and kept alive for an additional 10 minutes prior to PBS perfusion, agarose inflation, and lung harvest. For immunostaining, cleared tissues were incubated for two days with primary antibodies against α-SMA (1:1000, Abcam ab7817) or Ki67 (1:250, Fisher), followed by a two-day incubation with AlexaFluor-conjugated secondary antibodies (1:500, Invitrogen), with intervening PBS washes. Tissues were then sequentially stained with NHS-ester (1:3000, Invitrogen) and DAPI (0.1 mg/mL). Volumetric tissue imaging was then performed using a point-scanning confocal microscope (Zeiss LSM800) after the tissue was equilibrated in CUBIC-R^+^M imaging medium.

### Plasma analysis of secreted factors

Harvested blood was immediately centrifuged at 2,000 x g for 15 minutes at 4°C. For each time point, plasma from three male mice was pooled in equal volumes (100 µL total). Angiogenesis-related protein levels were assessed using the Proteome Profiler Mouse Angiogenesis Array Kit (R&D Systems), with immunoblots quantified using a custom MATLAB script.

### Quantitative image analysis

Images were processed using AIVIA or custom scripts written in MATLAB. In AIVIA, pixel classifiers were trained for each fluorescent signal to generate segmented channels for specific cell types. These channels were used to mask the DAPI channel to extract cell-type-specific nuclei, enabling quantification of cell number and volume via 3D analysis recipes. For K-means clustering, binary tile-scan channels (αSMA, GFP, DAPI) were merged into an RGB image. This image was binned to a resolution reflecting the high-magnification field of view (∼100×100 µm) and used as input for MATLAB’s k-means algorithm to cluster regions based on RGB intensity.

### K-Means Clustering (KMC) analyses

Whole-lobe tile-scan images were processed using a custom MATLAB script. Three fluorescence channels — Tie2-GFP, α-SMA, and nuclei (DAPI) — were used as input features for clustering. Each channel was first spatially binned into 40 × 40 pixel (∼100 × 100 µm) blocks by mean intensity to match high magnification image size and account for airspace in healthy lung feature while preserving tissue-level spatial information. Regions with imaging artifacts were manually excluded prior to binning. The three binned maps were then merged into a single RGB composite (R = α-SMA, G = Tie2-GFP, B = DAPI), and unsupervised K-means segmentation was performed using MATLAB’s *imsegkmeans*. The optimal cluster number was determined by the Elbow method; K = 3 (excluding empty space) was selected for bleomycin-treated lungs and K = 2 for uninjured controls. Clusters were assigned to biological categories based on centroid intensity rankings and confirmed by visual inspection. A fourth channel, the endothelial activation marker, was not included in clustering but was quantified per cluster region as a readout of endothelial activation. Cluster centroids were tracked across time points and projected onto the Tie2-GFP vs. α-SMA plane to visualize temporal trajectories.

## Statistical analyses

Statistical significance was determined by one- or two-way ANOVA or a two-sided Student’s t-test where appropriate (α=0.05). Outliers, defined as values >1.5x the interquartile range from the median, were excluded. Sample size (n) is indicated in the corresponding figure legends, and all data are presented as mean ± standard deviation.

