## supplementary data for "A 3D image atlas chronicling cellular and structural dynamics following lung injury identifies the aberrant expansion of endothelial cells that fail to form perfused vasculature"

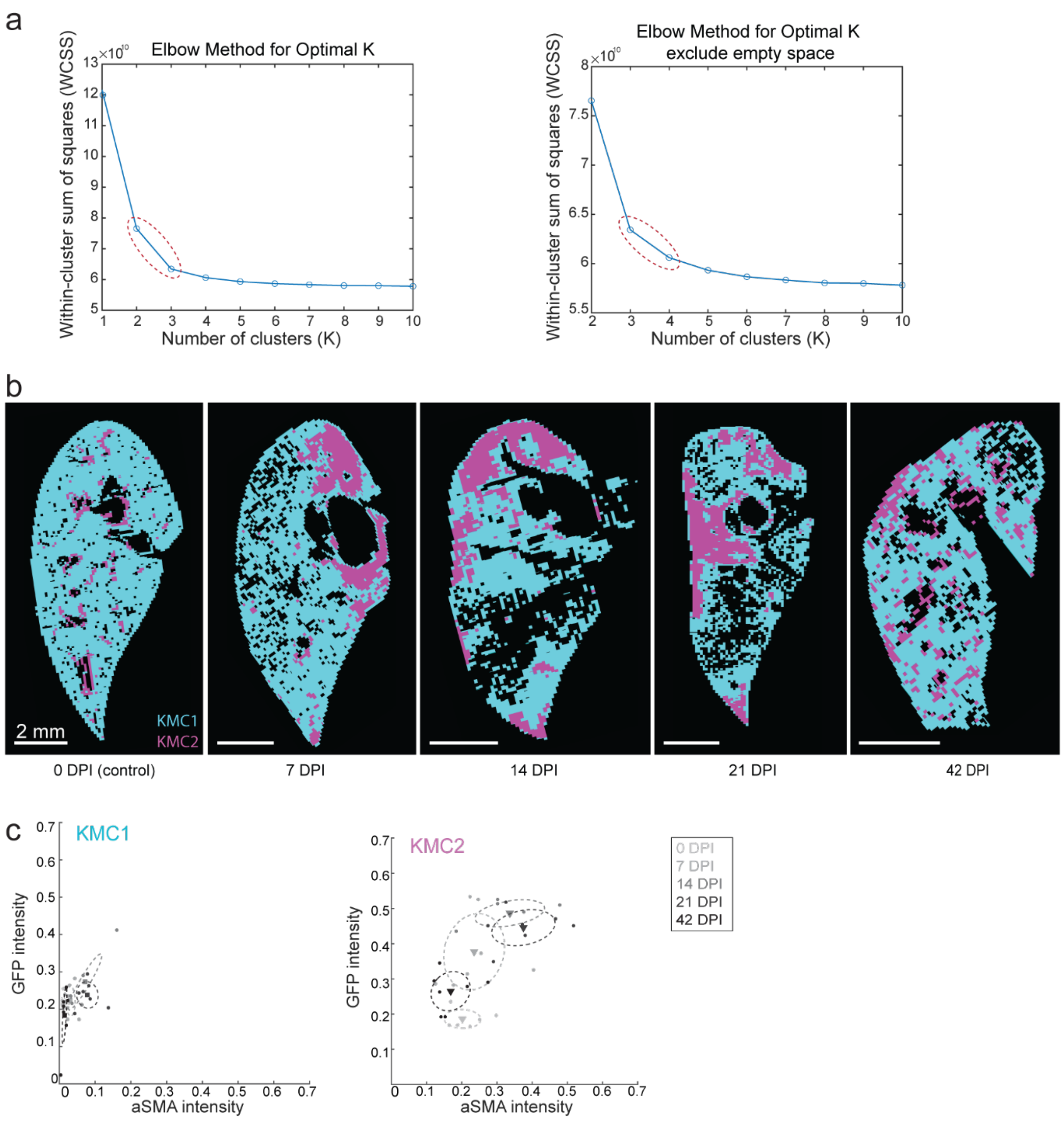


**Supplementary Figure 1: Determination of optimal cluster number (K) for fibrotic lung tissue: (a)** Elbow plot of the within-cluster sum of squares (WCSS) for K=1 to K=10. The distinct "elbow" at K=3 indicates the primary separation is between blank space, healthy and diseased tissue (K=2 for tissue, excluding empty space). **(b)** Representative K-means segmentation using K=2 (Cyan=Healthy, Magenta=Diseased). While this effectively isolates the lesion, it fails to capture heterogeneity within the injured area. **(c)** Scatter plot of the "Diseased" cluster from the K=2 model plotted in feature space. The large spread and distinct grouping within the magenta cluster (arrows) reveal underlying heterogeneity, justifying the refinement to K=3 to separately identify the endothelial-rich and myofibroblast-rich sub-regions. N=3 mice per time point.

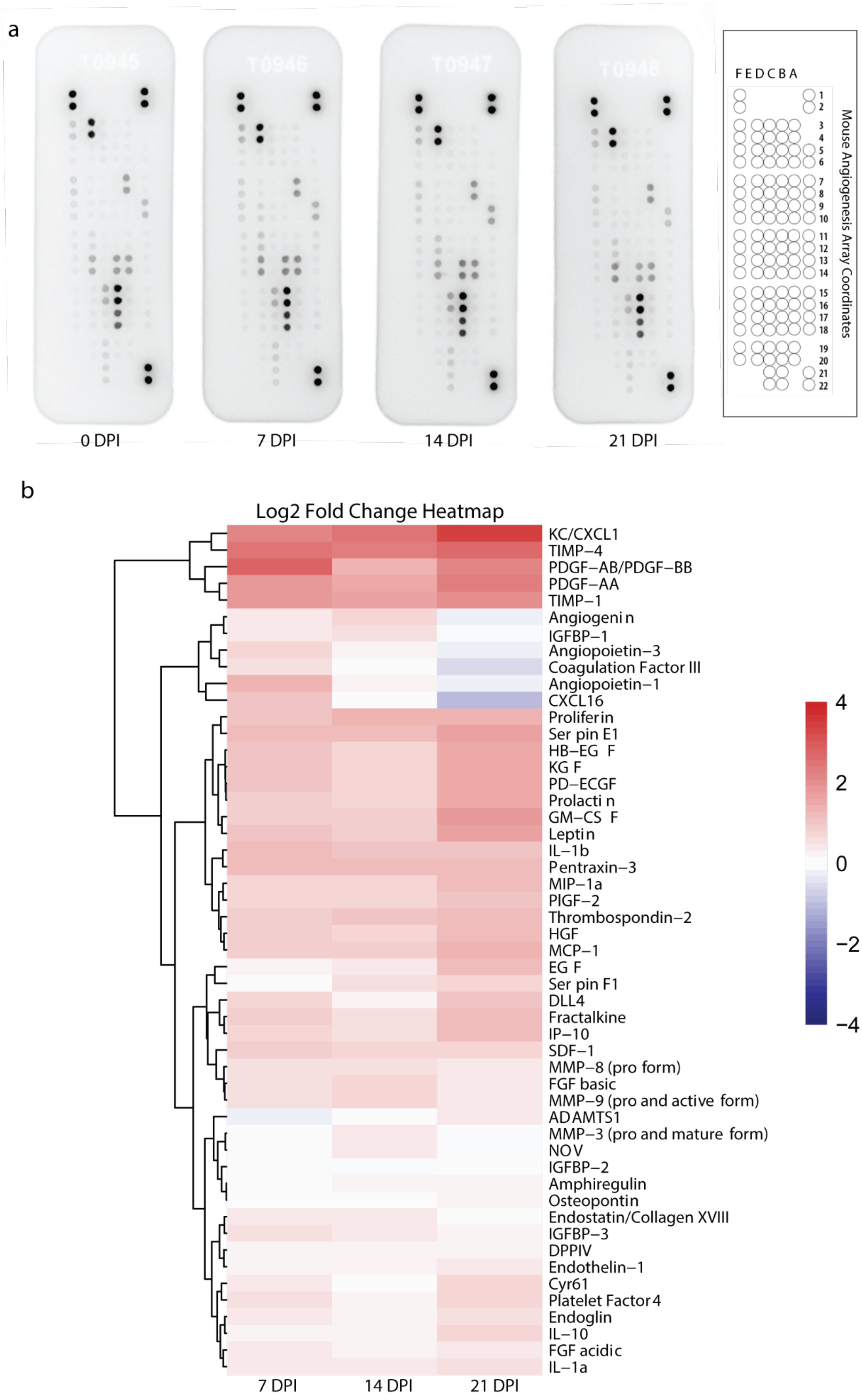


**Supplementary Figure 2: Upregulation of angiogenic factors precedes fibrotic remodeling:** (a) Representative images of the Proteome Profiler Mouse Angiogenesis Array membranes incubated with pooled mouse blood serum from 0 DPI (control) to 21 DPI post-injury. Each pair of dots represents a specific capture antibody in duplicate. N>=5 mice per time point. **(b)** Heatmap of the log2 fold change in pixel intensity relative to 0 DPI controls. The analysis reveals a broad upregulation of pro-angiogenic factors (e.g., TIMP-1, PDGF-AA, Endoglin) initiating at 7 DPI and persisting through 42 DPI, providing a molecular basis for the observed endothelial expansion.

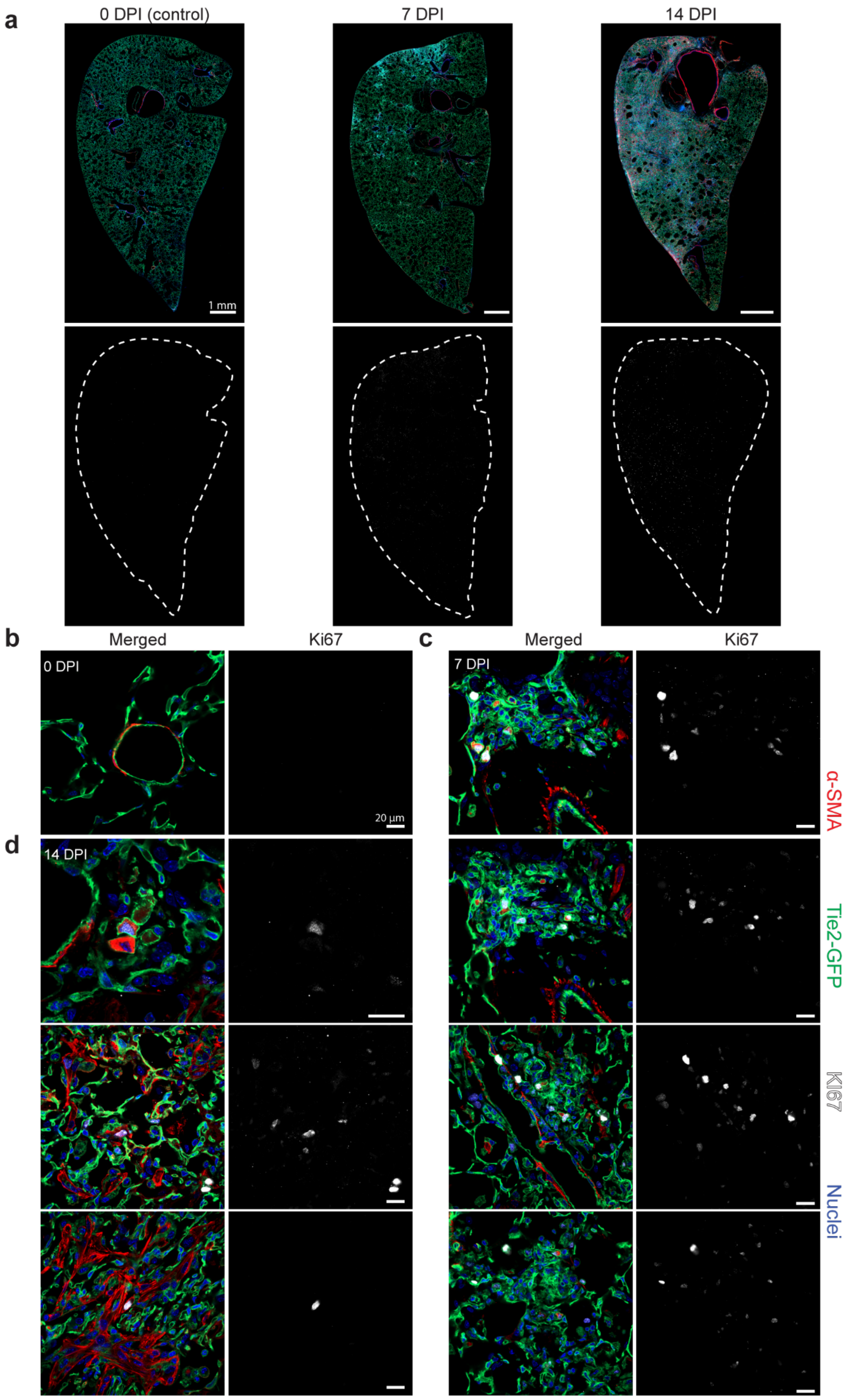


**Supplementary Figure 3: Spatiotemporal analysis of cell proliferation during fibrosis progression:** **(a)** Representative whole-lobe tile scans showing the distribution of proliferating cells (Ki67, white) relative to vasculature (Tie2-GFP, green) and MF (α-SMA, red). Note the marked increase in proliferation at Week 1 compared to the quiescent Week 0 baseline. Due to the sparse, single-cell nature of the signal, the lung boundary is outlined (dashed line). **(b)** High-magnification view of uninjured lung (Week 0) showing healthy vasculature with minimal to no Ki67 expression. **(c)** At Week 1, proliferation is prominent within the GFP+ EC and forming KMC3-like regions. Note the Ki67+ nuclei within GFP+ ECs and dual-positive transition cells. **(d)** At Week 2, while Ki67 signal attenuates in ECs and dual-positive cells, Ki67+ nuclei are frequently observed within α-SMA + MFs (bottom row). This indicates KMC2 fibrotic core expansion is driven in part by the local proliferation of MFs. N=3 mice per time point and n>2 tile scans per mouse.

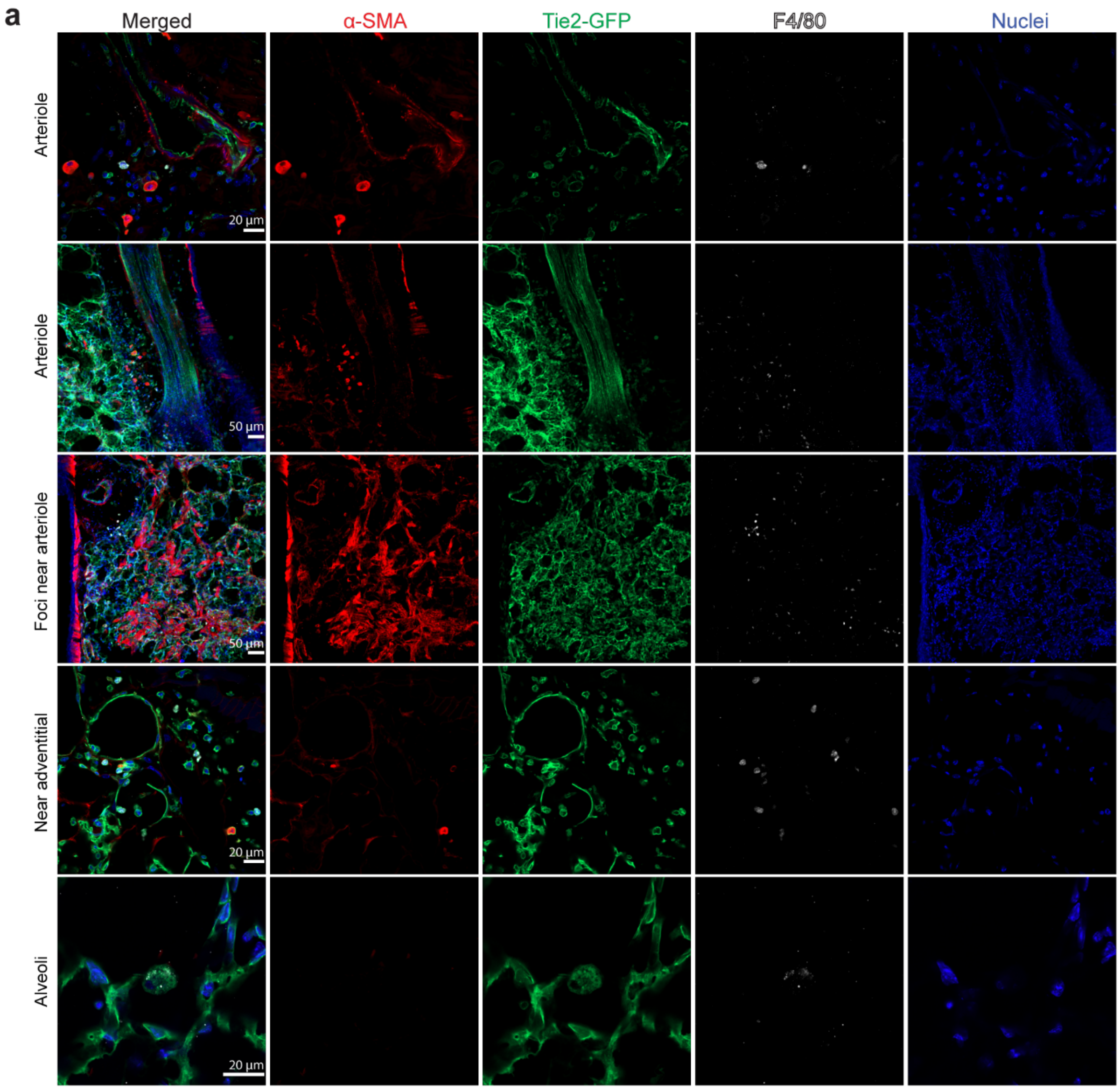


**Supplementary Figure 4: F4/80 staining distinguishes macrophages from the expanded endothelial network: (a)** Representative high-resolution confocal images of injured lung tissue co-stained for α-SMA (red), Tie2-GFP (green), and the macrophage marker F4/80 (white). F4/80+ cells appear as discrete, single cells primarily located in the perivascular adventitia or airspace. They are morphologically distinct from the dense, interconnected Tie2-GFP+ networks and do not colocalize with the GFP signal in the parenchymal injury zones, confirming F4/80+ macrophages are not a major source of the expanded GFP+ cell population.

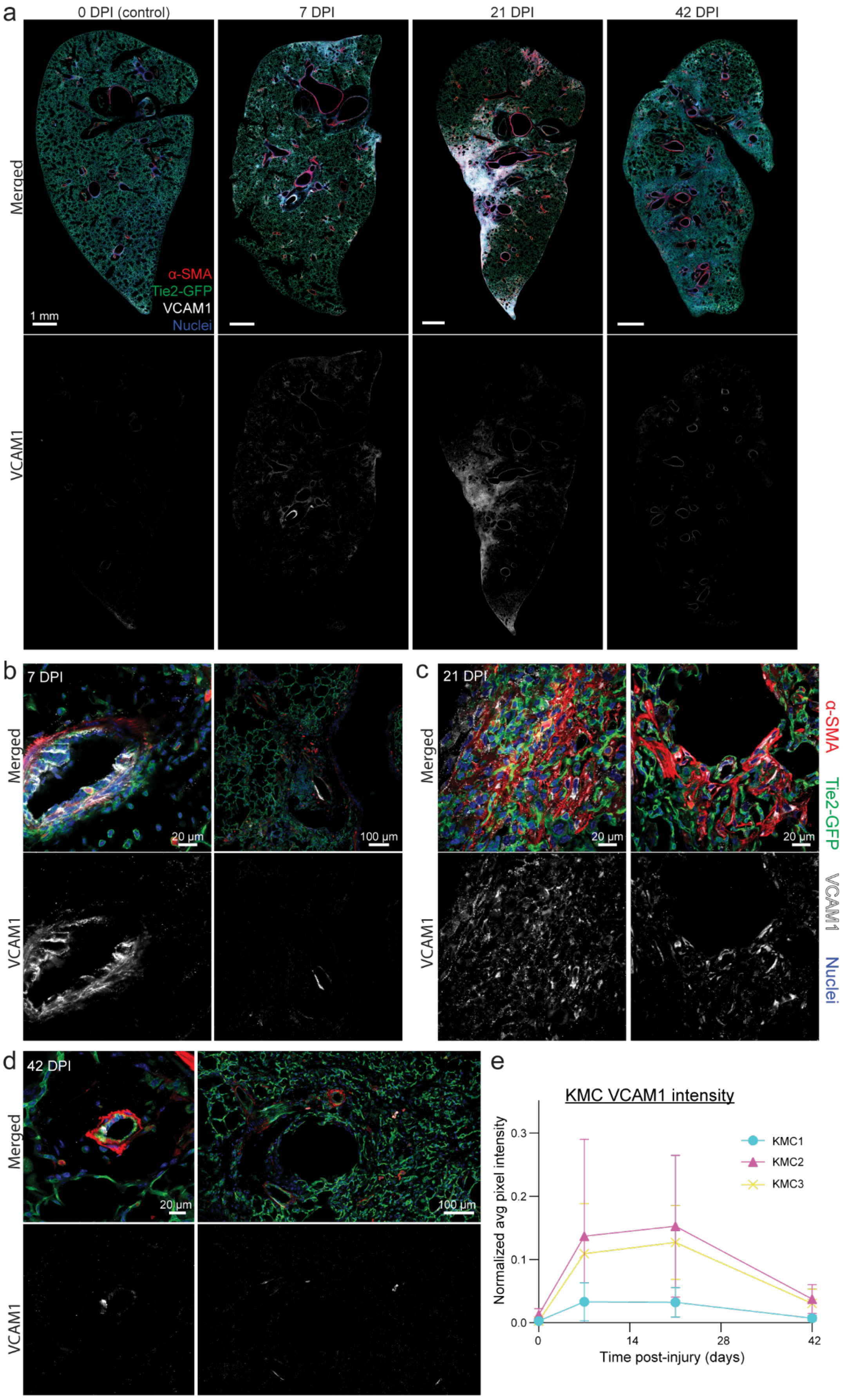


**Supplementary Figure 5: Spatiotemporal dynamics of VCAM-1 expression:** (a) Whole-lobe tile scans showing VCAM-1 expression (white) relative to the fibrotic injury (Tie2-GFP/ α-SMA). VCAM-1 signal peaks transiently at Week 1, specifically tracing large vascular structures, before diminishing by Week 3. (b-d) High-resolution imaging of VCAM-1 expression. (b) Week 1: The most intense VCAM-1 staining is localized to the endothelium of large vessels. Note the hypertrophic, cuboidal morphology of the VCAM-1+ ECs and their protrusion into the lumen, indicative of acute activation. (c) Week 3: VCAM-1 signal shifts from a distinct vascular pattern to diffuse, lower-intensity signals within the fibrotic core, suggesting waning EC activation and low-level expression by activated mesenchymal cells. (d) Week 6: VCAM-1 signal returns to near-baseline levels in the alveolar space, with occasional signal on the inner wall of large vessels. (e) Quantification of normalized VCAM-1 pixel intensity. Expression peaks sharply at Week 1 in both diseased regions (KMC2 and KMC3) before declining, consistent with an acute inflammatory phase. Data are mean ± s.d. N=3 mice per time point and n>2 tile scans per mouse.

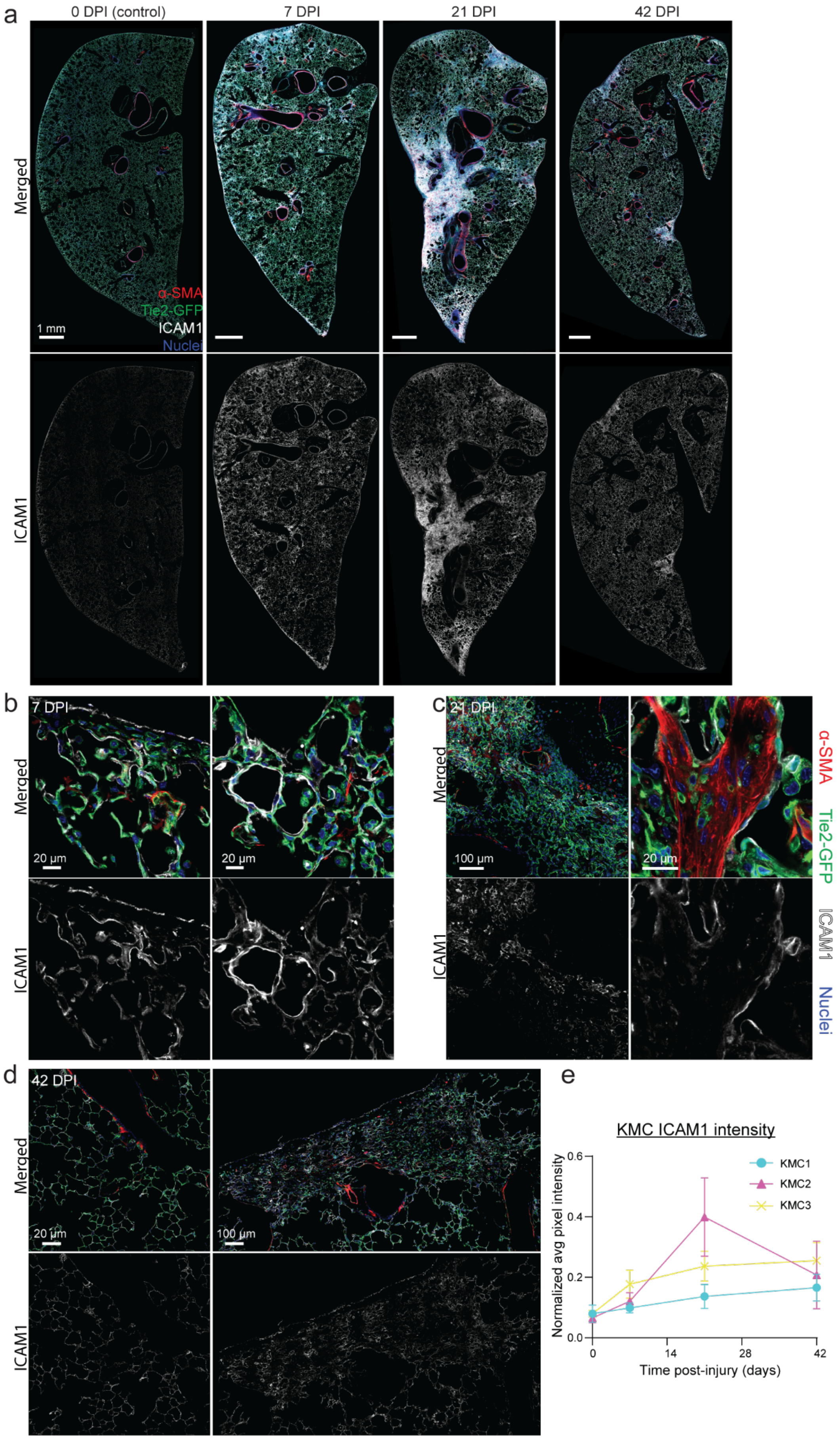


**Supplementary Figure 6: ICAM-1 expression defines a persistent, pan-pulmonary inflammatory state.** (a) Whole-lobe tile scans of ICAM-1 expression (white). Unlike VCAM-1, ICAM-1 signal increases globally at Week 1 and progressively intensifies, peaking at Week 3 within the fibrotic regions. (b-d) High-resolution imaging of ICAM-1 distribution. (b) Week 1: Global upregulation of ICAM-1 across both healthy and injured regions, highlighting a systemic response to injury cytokines. (c) Week 3: Maximal ICAM-1 expression. Note intense signal throughout the dense KMC2 core, indicating contributions from both the expanded endothelium and injured epithelium/myofibroblasts. (d) Week 6: ICAM-1 levels remain elevated compared to controls, indicating a "molecular memory" of inflammation. (e) Quantification of normalized ICAM-1 pixel intensity. ICAM-1 shows a delayed peak at Week 3, with the highest intensity in the KMC2 fibrotic core (Magenta), supporting a mixed cellular source (EC + Epithelial + MF). Data are mean ± s.d. N=3 mice per time point and n>2 tile scans per mouse.

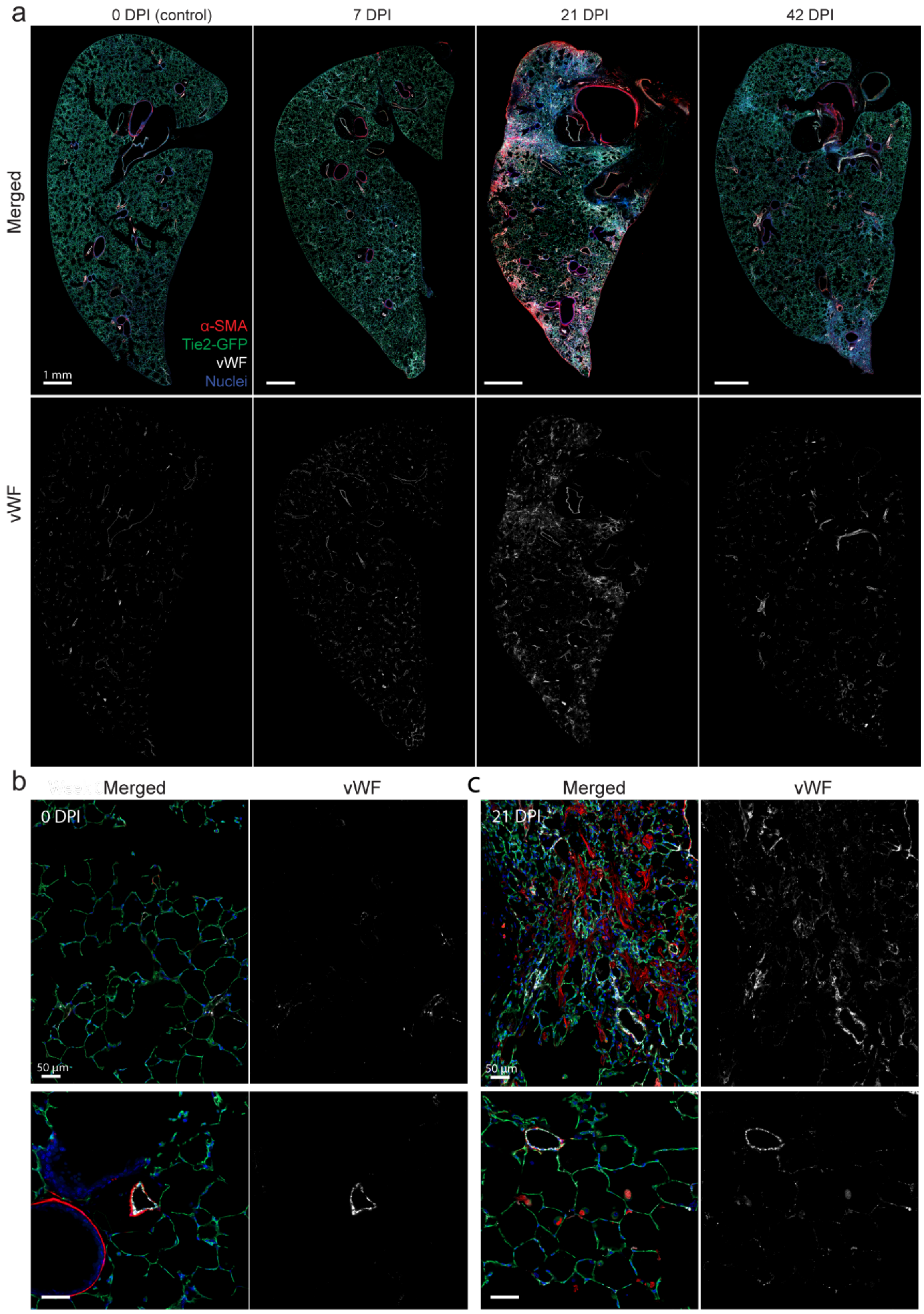


**Supplementary Figure 7: Abnormal expansion of vWF into the distal capillary network:** (a) Whole-lobe tile scans showing vWF (white) expression progression. At Week 0, signal is sparse and restricted to large vasculature. By Week 3, intense vWF signal extends throughout the parenchymal injury zones before resolving at Week 6. (b) High-resolution imaging of Week 0 (Baseline) showing vWF restriction to large arterioles/venules (bottom left), while the alveolar capillary network remains vWF-negative. (c) High-resolution imaging of Week 3. Note the aberrant, high-intensity vWF expression extending into the dense, disorganized capillary networks within the injury zone (merged panel), confirming endothelial identity of the expanded KMC3 region. N=3 mice per time point and n>2 tile scans per mouse.

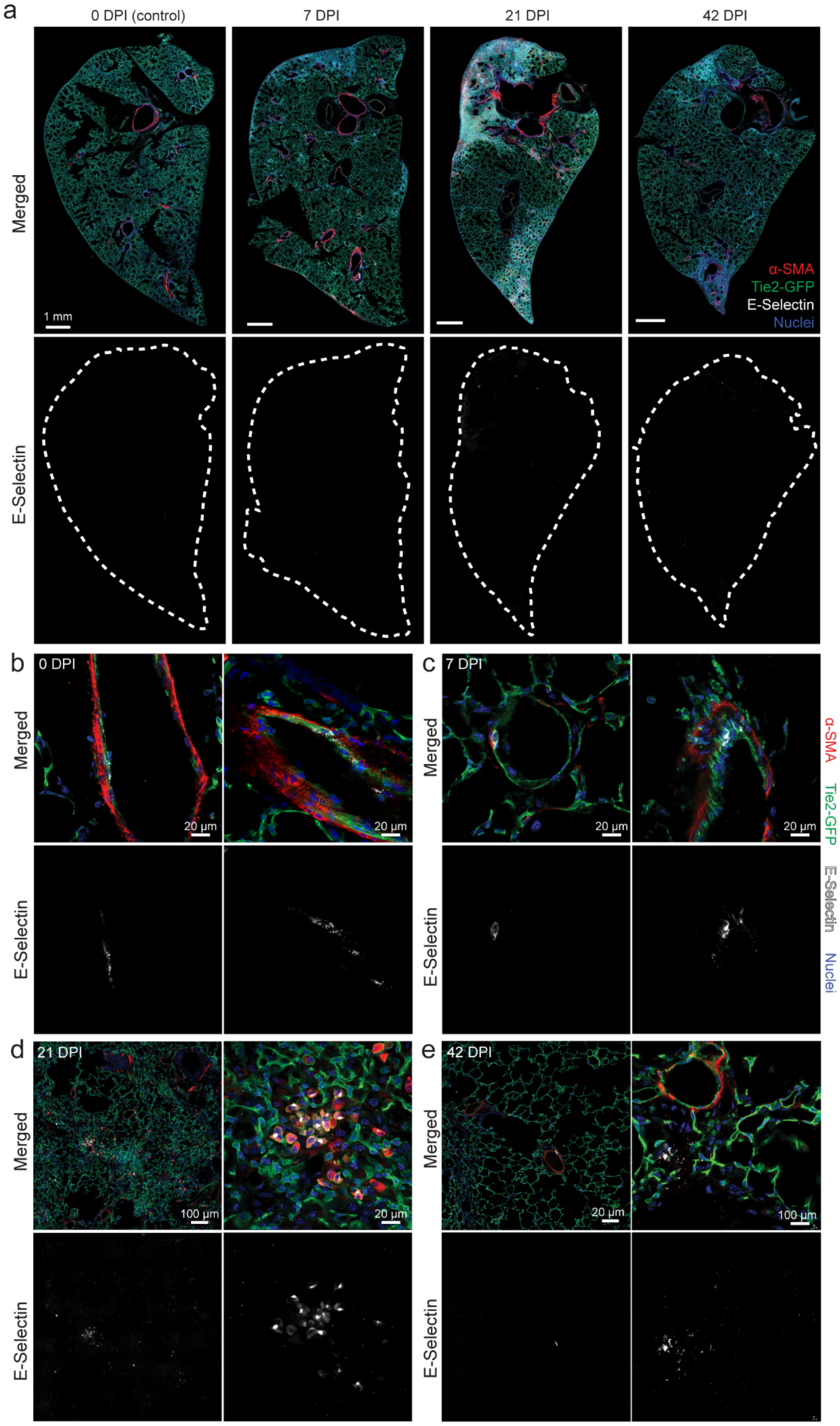


**Supplementary Figure 8: E-selectin expression identifies activated and transitioning endothelial cells.** (a) Whole-lobe tile scans of E-selectin expression. Due to the sparse, single-cell nature of the signal, the lung boundary is outlined (dashed line). Unlike global inflammatory markers, E-selectin remains spatially discrete throughout the time course. (b-e) High-resolution imaging of E-selectin dynamics. (b) Week 0: Minimal baseline expression restricted to rare ECs on the inner wall of large vessels. (c) Week 1: Upregulation of E-selectin on the inner wall of large vessels, often clustering at sites of immune cell interaction (merged panel). (d) Week 3: Maximal E-selectin expression is identified on dual-positive cells (GFP+/α-SMA +, arrows). The specific localization of this endothelial adhesion molecule to cells expressing mesenchymal markers strongly supports an endothelial-to-mesenchymal transition (EndoMT)-like phenotype. (e) Week 6: Resolution phase. Coherent vascular staining is replaced by fragmented, punctate signals that do not co-localize with cells, consistent with endothelial apoptosis and proteolytic shedding of E-selectin during vascular regression. N=3 mice per time point and n>2 tile scans per mouse.

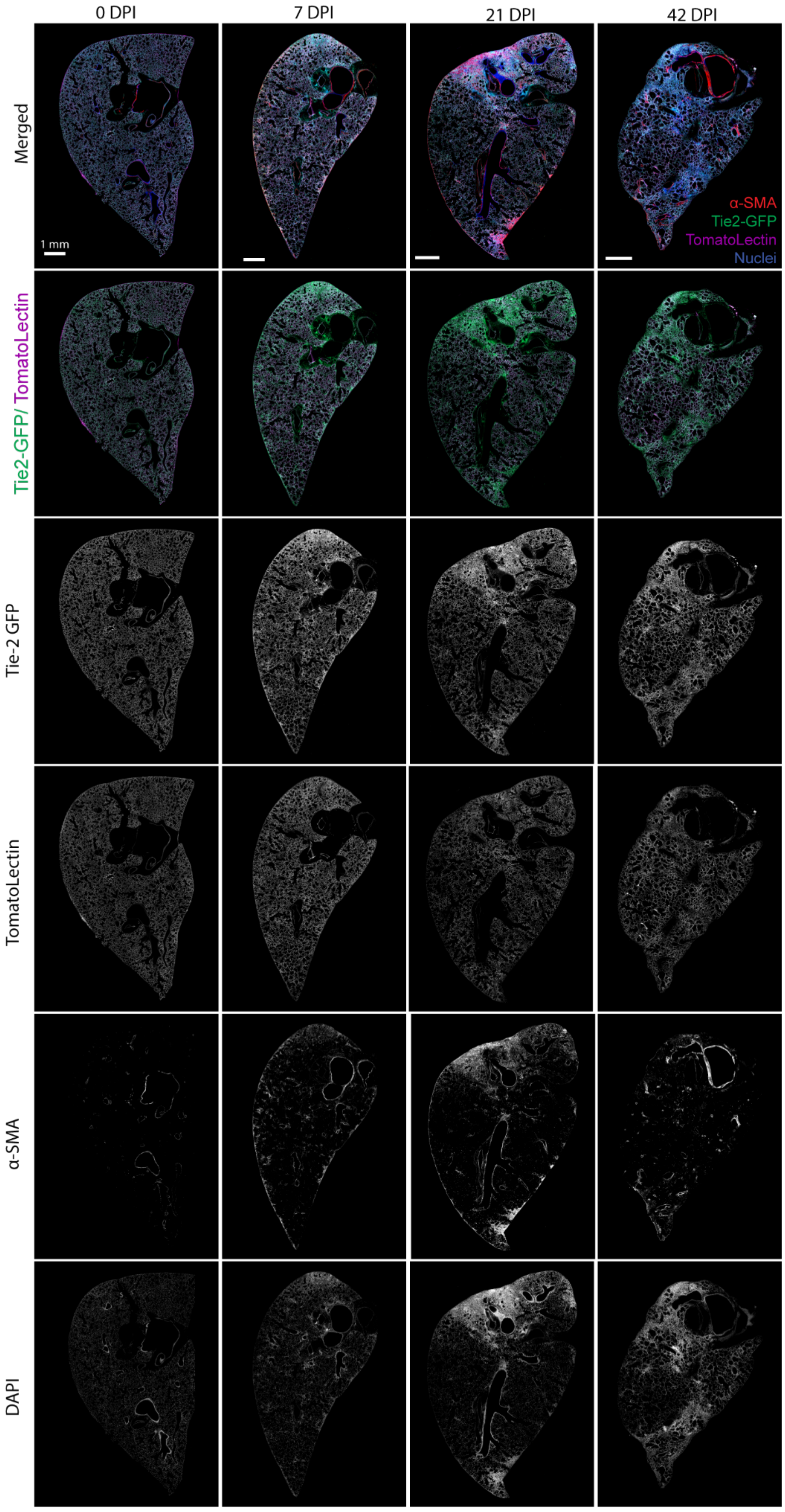


**Supplementary Figure 9: Whole-lobe visualization of perfusion deficits:** Representative whole-lobe tile scans of Lectin perfusion (purple) overlaid with the EC lineage trace (Tie2-GFP, green) and MFs (α-SMA +, red) across the time course. Rows: (1) Merged composite; (2) Tie2-GFP lineage trace showing total endothelium; (3) Tomato Lectin showing perfused vasculature; (4) α-SMA showing MF and smooth muscle cells; (5) DAPI (nuclei). Note the significant mismatch between the total endothelial area (Row 2) and the perfused area (Row 3) in the injury zones at Week 1 and 3, visually confirming the "perfusion gap" quantified in Figure 5. N=3 mice per time point.
